# Secondary structure of pre-mRNA introns for genes in the 15q11-12 locus. Mapping of functionally significant motives for RNA-binding proteins and nucleosome positioning signals

**DOI:** 10.1101/409243

**Authors:** Viya B. Fedoseyeva, Irina A. Zharinova, Alexander A. Alexandrov

**Author notes:** Corresponding author (VBF). 1st set of equal contributors. These authors contributed equally to this work (Irina A. Zharinova, Alexander A. Alexandrov).

## Abstract

In this study, we identified reproducible substructures in the folded structures of long intron RNAs for recursive spliced variants and annotated pre-mRNA for *GABRB3 and GABRA5*. We mapped the RNA motives recognized by RNA-binding proteins for the specified locus and characterized the area of preferred localization. A comparison of pre-mRNA variants revealed the dominant type of protein potential effects. We determined the structural specifics of RNA in the dense Alu cluster and clarified the analogy of apical substructure to the A-Xist fragment of transcriptional variant. Mapping of the nucleosome potential reveals alternation of strong and weak signals at the 3’-end portion of *GABRB3* and clusters of nucleosome positioning signal in the vicinity of the Alu cluster. Distribution of simple oligonucleotides among reproducible substructures revealed an enrichment in Py-tracts; for some of them, this may be considered as a complementary supplement to the Pu-tract enrichment of ncRNA Malat1 as a component of nuclear speckles. The secondary structure elements of bidirectional transcripts are predisposed for somatic homolog pairing in this locus, as was previously shown experimentally.

A model of potential intron RNA influence on splicing has been suggested based on its interaction with Py-tract-binding RNP, serine-arginine *SRSF* proteins, ncRNA Malat1, as well as the action of Alu cluster.

## Introduction

The splicing model for exons surrounded by long introns is based on the assumption of a pre-assembly of future spliceosome elements [1]. Splicing processes are assisted by components of other processes, such as transcription (RNA-Pol II CTD) and chromosomal activation [2-6], including the SAGA and SWI/SNF complexes [7-12]. A large fraction of introns spanning thousands of nucleotides participates in co-transcriptional splicing (coTS) without hindering splicing [13]; however, among them, there is a fraction (up to 20%) that may be subject to splicing at the post-transcriptional level (postTS) [14,15]. For example, the first long intron(s) is removed from pre-mRNA more slowly than the others [14] and thus is the first candidate for postTS. The role of the large introns themselves, at least of their main portion, remains poorly understood in the splicing process. Still, a separate facet of the interaction has long been known, namely, that the association b**e**tween the nascent RNA and splicing factors in the nucleus is intron-dependent [16]. The significance of long introns is emphasized by the phenomenon of protecting long pre-mRNAs from premature cleavage and polyadenylation [17-18]. The role of long introns can be clarified by identifying the interaction of pre-mRNA (annotated as coding and/or *in silico* predicted RNA) with RNA-binding proteins and other RNAs, for example, non-coding RNA, pertaining to spliceosome pre-assembly. Non-coding RNA Malat1 is associated with recruitment of SR family pre-mRNA-splicing factors from nuclear speckles (NS) to the transcription sites [19]. In addition, it is also known that the splicing of exogenous pre-mRNAs occurs when they recruit the inter-chromosomal granular cluster (GC), e.g., serine-arginine NS cluster [20, 21]. The composition of the granules includes *SRSF1, SRSF2, U2snRNP* proteins, MALAT1 and other components of the spliceosome. The peri-chromatin filament [PF] region containing the endogenous nascent RNA and associated proteins also can recruit GC granules [20-23], although the interaction of PF and GC is not always obvious [24,25]. Splicing strongly depends on the presence of pyrimidine tracts in RNA [20, 26-27]; their removal leads to attenuation of splicing and granule binding. The role of pyrimidine tracts pertains not only to the nearest branch site but also to some cryptic sites.

This work aimed to identify the RNA features at protein binding sites in the case of one-and two-dimensional presentation. This involves determination of the RNA secondary structure in long introns. First, we focus on proteins involved in the composition of NS as well as the proteins interacting with pyrimidine oligomers. In addition, since according to the model, the nucleosome formation potential influences coTS, we elucidated the peculiarities of nucleosome and *CTCF* mapping at the DNA level because strong nucleosome signals and *CTCF* [28] may influence transcription pauses. The current work examined the DNA locus 15q11-13. The documented phenomenon of somatic pairing of homologous chromosomes was also included in our consideration as pairing disorders and emergence of diseases often occur simultaneously.

## Features of the 15q11-13 locus

This locus encodes the α5, β3, and γ3 genes of GABAA receptor subunits. They do not belong to the group of most-common receptors subunit genes (α1,2, β1,2, γ1,2, etc.), but a wider range of structural and functional properties of α5(*GABRA5*) and β3(*GABRB3*) genes and their association with neurodegenerative diseases make them the most attractive for sequence analysis. The β3 gene is expressed not only in the brain but at lower levels in other tissues; as a part of locus, it participates in somatic homologue pairing in late S phase [29] in lymphocytes as well as in neuronal tissue and in *in vitro* systems [30,31]. The β3 gene is the shortest among β genes, although other *in silico* predicted bi-directional transcript variants as well as recursive splicing variants have been suggested in addition to experimentally annotated ones. The bi-directionality is supported by the presence of spliced and unspliced EST sequences in the *antisense* and *sense* versions. Elucidation of the structural and functional properties of the bi-directionally oriented Alu repeat cluster at the beginning of β3 gene is a separate problem, and despite the knowledge of multiple properties of this repeat type [32], the desirable completeness of information is not achieved, especially at the transcriptional level.

The multiple annotated variants of pre-mRNA and protein isoforms for the β3, α5 genes are significantly different in size and stages of expression (foetal and adult). Expression of a long variant 1,2 with a long intron at the beginning of the transcript (after the cassette exons) is associated in the brain with a foetal developmental stage, the intermediate length pre-mRNA (variant 3) is also expressed in brain and at lower levels in lungs, cardiomyocytes, and germinal cells [33]. The shortest pre-mRNA variant 4 (core-part) is expressed in adult brain. The β3 gene is of importance due to its association with the Angelman syndrome [34-38], with the Prader-Willi syndrome (multiple deletions of a significant middle portion of a long intron) [39], with epilepsy (point mutations), and autism [40]. *GABRB3, GABRA5* are considered as candidate genes responsible for panic disorder [41].

## Materials and Methods

We used the calculation of minimum free energy (MFE) and RNA folding by Markham and Zuker’s programme UNAFOLD (42). Computations were supported by MS Windows (32-bit architecture), Linux (64-bit), Cluster/img.ras.ru/galaxy resource and MFOLD [43] for thermo-dynamic parameters determination, as well as by Microsoft Office, CorelDraw and Delphy7 for mapping of nucleotide motives. For calculations of nucleosome positioning [NP] potential we used a previously written Turbo Pascal script [44]. For the primary sequences and annotated variants of pre-mRNA, *in silico* prediction by Genscan and Gene id programmes was done for promoters (annotated and hypothetical), EST, GC-rich regions, and Alu repeats using https://ncbi.nlm.nih.gov (GenBank) and https://genome.ucsc.edu as data sources.

## Results and discussion

This study is divided into several parts: (a) two folding methods for large intronic RNAs, (b) mapping of recognition sites for RNA-binding proteins at the primary nucleotide sequence level and at the level of secondary structures in intronic RNA, (c) evaluation of oligonucleotide occurrence close to the motives recognized by RNA-binding proteins and their complementary counterparts, for the folded secondary structure branches (set of reproducing helix-loop structures) and intermediate fragments, (d) examination of RNA folding in the dense Alu cluster and A-Xist–like apical substructure associated with it, (e) the DNA level mapping of nucleosome potential (NP) and proposed transcriptional pauses associated with CTCF binding [28], and (f) analysis of communication between homologous chromosomes mediated by *antisense* and *sense* pre-mRNA with the aim of offering a possible mechanism to explain the initiation of pairing between homologous chromosomes.

## Scheme of 15q11-12 locus

Fig. 1A, B shows localization of the Alu repeats in two orientations; the clusters of them can be seen at the beginning (5’ end) of intron 3 (GenBank, *GABRB3* gene) and at the end of intron 3. One can see experimentally identified transcript variants, corresponding to different protein isoforms for *GABRB3* and *GABRA5* genes (Fig. 1C-L), the computationally predicted transcripts matching the experimental ones, sites of recursive splicing for long introns (Fig. 1N-S), promoters, GC-rich region (Fig. 1M, O) and EST including un-spliced for both orientations (hg38 assembly) (Fig. 1T,U).

The mapping data for sites along the nucleotide sequence of locus 15q11-12 are shown in Fig. 2 for RNA-binding proteins: for serine–arginine *SRSF1* (aliases *ASF, SRp30a, SFRS1, SF2*), *SRSF2* (aliases *SC35, SFRS2, SRp30b, SFRS2A*), *SRSF5* (aliases *SFRS5, SRP40, HRS*), *hnRNP A, C* and *PTB* proteins. The data are based on functional methods, immunoprecipitation and SELEX. Since *GABRB3* and *GABRA5* genes have different orientations, the mapping was carried out for the (+) strands and (–) strands in accordance with gene orientation.

**Fig 1.**
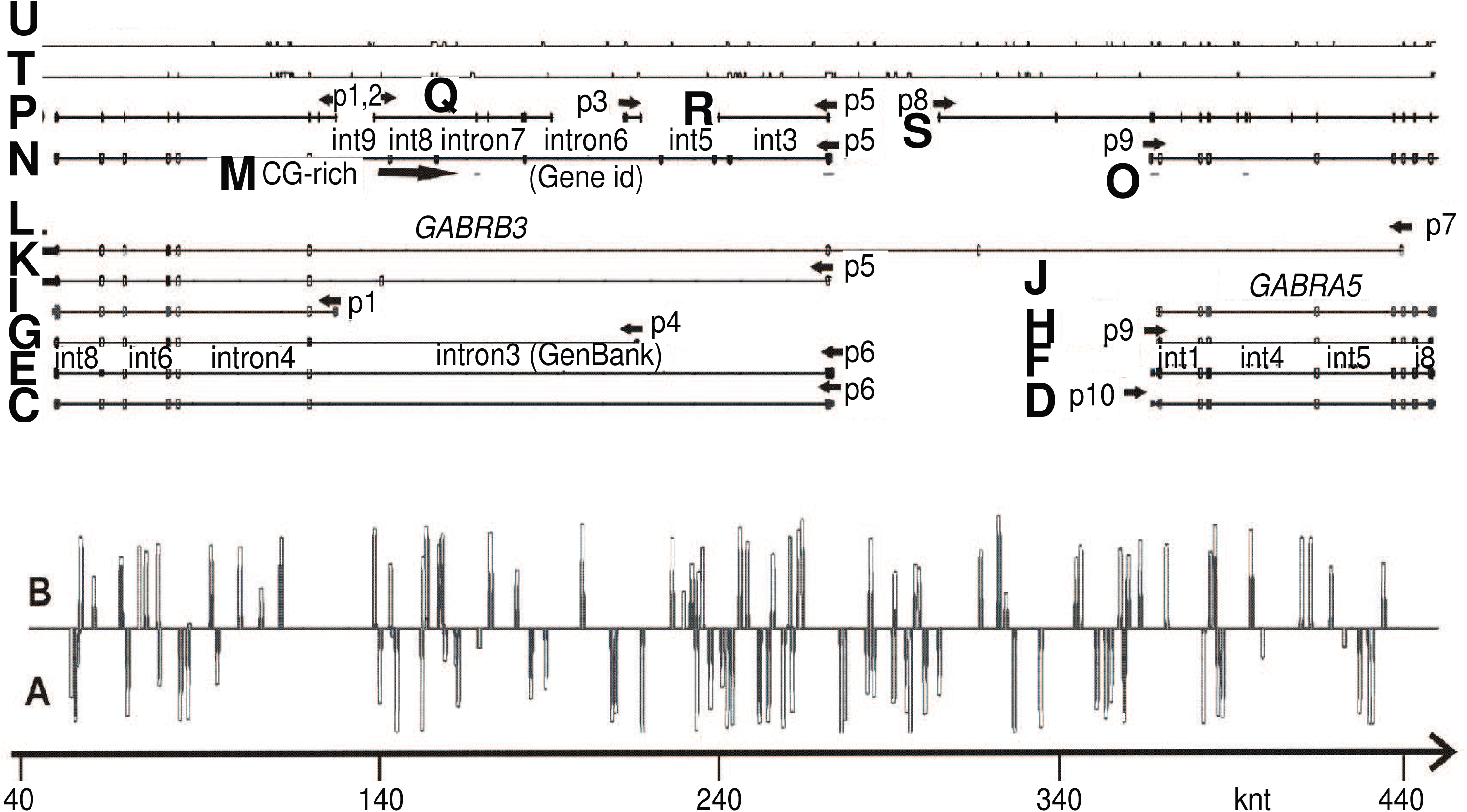
Scheme of *GABRB3* and *GABRA5* operon, annotated and *in silico* predicted transcription variants, EST and Alu repeats. (A), (B) – Alu repeat localization in the (+) and (-) orientation. (C) KJ534842, var2, adult brain. (D) L08485 (adult brain). (E) AK315311, var 2 (foetal brain). (F) BC113422 (brain and lung). (G) AK302822, var3 (brain, cardiomyocyte, lung, testis *et al*). (H) BC111979 (brain and lung). (I) AK295167, var4. (J) BC011403 (retinoblastoma). (K) BC010641, var1 (retinoblastoma). (L) CR749803 (retina). (M) GC-rich regions. (N-O) Gene id (-). (P) Genescan (-). (Q) Genescan (+). (R) Genescan(-). (S) Genescan (+). (T) EST (-). (U) EST (+). Start point (40 knt) corresponds to 26540 knt for chr15 (hg38 assembly). (C) -(L) Annotated human mRNA from GenBank. (N)-(O) *in silico* predicted mRNA by Genscan, Gene id programmes. Annotated promoters P1, P4-6, 9, 10. (C) -(L) GenBank mRNA variants, http://genome.ucsc.edu

**Fig 2.**
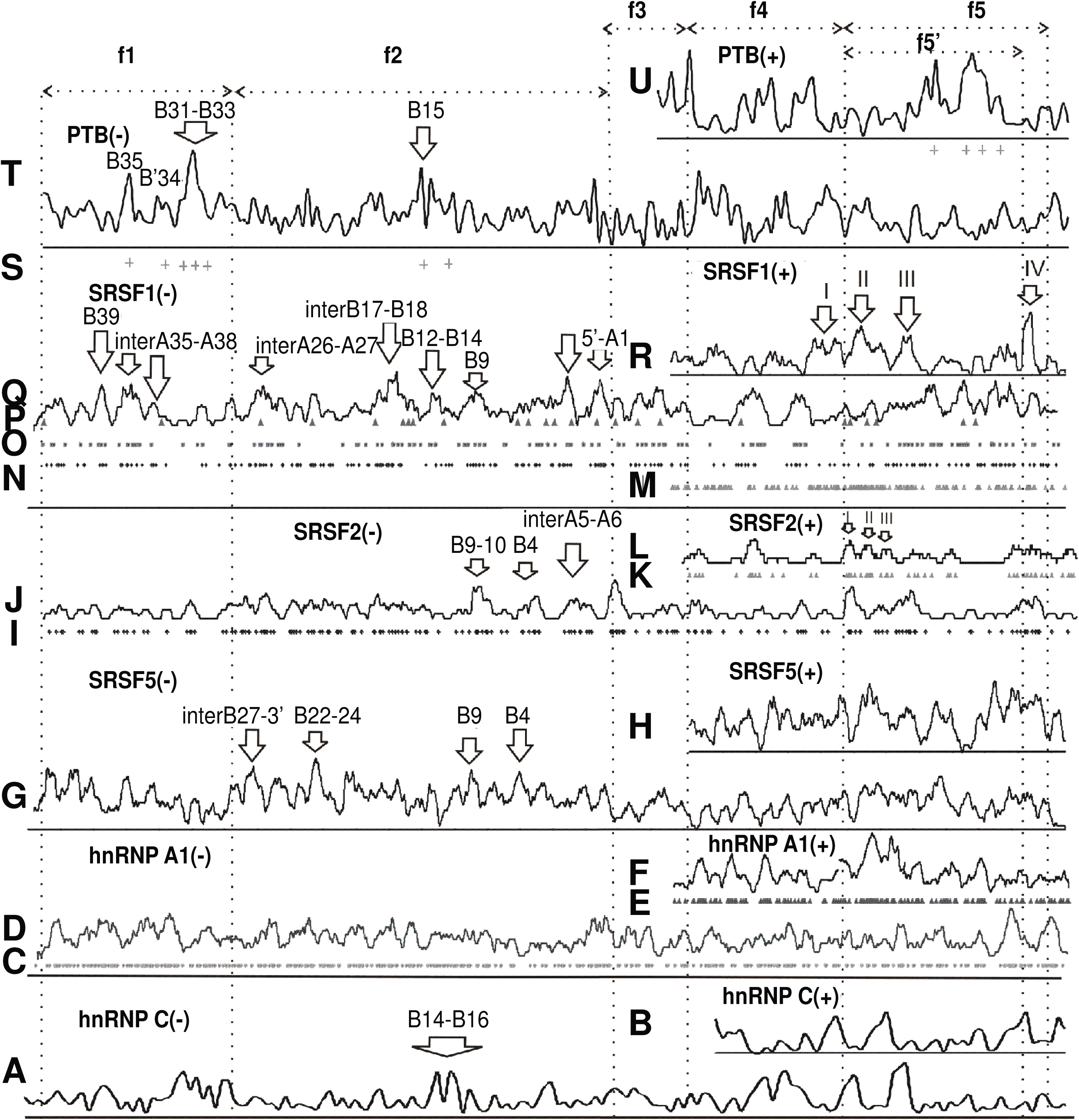
Mapping of sites for RNA-binding proteins. (A) *hnRNP C* site (complement to 5T (-)) [45] after averaging. (B) *hnRNP C* binding site (motif 5T (+)) after averaging. (C) *hnRNP A1* site (complement to TAGGGA/T (-)) [46]. (D). Averaged data from (C). (E) *hnRNP A1* site (motif TAGGGA/T (+)). (F) Averaged data from (C). (G) *SRSF5* site (complement to CDGCA (-)) [47]. (H) *SRSF5* site (motif CDGCA (+)). (I) *SRSF2* site (complement to AGGAGAT and GRYYCSYR (-)) [48, 49]. (J) Averaged data from (I). (K) *SRSF2* site (motif AGGAGAT and GRYYCSYR (+)). (L) Averaged data from (K). (M) S*RSF1* SRSASGA, RGAAGARR, RGAAGAAC sites (+). (N) *SRSF1* sites (complement to SRSASGA (-)) [47]. (O) *SRSF1* sites (complement to RGAAGARR(-)) [50]. (P) *SRSF1* sites (complement to RGAAGAAC (-)) [51]. (Q) Averaged data from (N-P). (R) Averaged data from (M). (S) *PTB P* motives as in (T) incorporated in Py-rich tract (> 15 nt) for (-). (T) *PTB P* sites (complement to TTCT, TCTT, CTCTCT (-)) after averaging. (U) Motives of *PTB P* sites (TTCT, TCTT, (C)TCTCT (+)) [52-54] after averaging. f1 fragment -core-part, f2 fragment – intron 3 (GenBank, *GABRB3*), f3 fragment – between P5 and P8 promoters, f3+f4 fragments – between *GABRB3* and *GABRA5* regions, f3+f4+f5’ fragment - two first introns for a long variant CR749803 (Fig. 1L), f5 fragment - *GABRA5* gene. (G) - (P) R-purine, Y-pyrimidine, S: G or C; D: A, G or U.

## Two folding methods for large intronic RNAs

First, in accordance with *in silico* predicted splicing sites (Gene id programme), we subdivided the longest intron of pre-mRNA (149 knt) (Table S2) into smaller fragments. Their lengths allow acceptable time needed for computation of secondary structure folding of intronic RNAs. These fragments may be considered as corresponding to recursive splicing. Fig. 3 depicts long intronic RNA of *GABRB3* gene that together with a core-part (Fig. 4) constitutes transcription variants 1,2. The core-part together with small exons/introns 1, 2 and 5’UTR corresponds to variant 4. Truncation of long 149-knt intron to 95 knt and joining it with the core-part gives rise to variant 3 (Fig. 5). The lost fragment incorporates the Alu cluster with a presumably important function that leads to an enhancement of expression in some tissues. Variants 1, 2 (bivalence is due to an alternative splicing of starting exons) are expressed in brain at a foetal developmental stage, whereas variant 3 is largely expressed in adult brain and, to a lesser extent, in cardiomyocytes, lung, testis and in muscles [Proteomics, GenBank]. Variant 4 predominantly expresses in adult brain. Additionally, there is a very long transcript expressed in retina. According to the latest data, the locus transcription is bi-allelic in brain, and in disease, it is partially biased to mono-allelic variants [31]. In Fig. 3, the short constitutive introns are numbered 1 and 2, recursive intron numbering runs from number 3 to 9 (Gene id), and their entire combination corresponds to the constitutive intron number 3 (GenBank). The exons are presented schematically without showing their secondary structure. Alu repeats are indicated by letters A with the occurrence number in double-stranded state. Numbering by letter B marks the branches that may be considered as spatially separated substructures consisting of rows of alternating loops and helices. The folding images for intronic RNAs correspond to thermodynamically optimal structures, whereas the suboptimal ones have minimal differences and are not considered in the context of fragments of such length. The coordinates of structural elements relative to the genomic sequence are given in Table S2 for the hg38 assembly of *Homo sapiens* genome (GenBank). These folding images are further used as the basis for mapping of RNA-binding proteins motives.

Second, the sliding window method was used for non-recursive folding variants, if such exist, to estimate the possibility of identification of folding peculiarities in long intronic RNAs, for example, intron 3 (GenBank, 149 knt) (Fig. 6). This non-recursive folding may be realized at the early interphase when splicing is delayed compared with transcription. For each sliding window used for folding, in the resulting structure we distinguish the branches as clusters of concentrated helix-loop chains. Some of them are reproducible substructures when the length and position of the sliding window vary. Some of them coincide with the branches of the same coordinates for *in silico* predicted fragments (recursive variant) of the same intron (Fig. 3). They are labeled with Bn as in previous description. The construction of the integral structure of the 149-knt intron (variant 3) is ambiguous due to the complexity and time required for calculation of the whole structure, but it is possible to link up randomly partitioned shorter nucleotide fragments (4-5 units) of reasonable length for minimization of total calculation time. After multiple attempts of composing, we selected those which have maximal numbers of reproducible branches. One of these potential variants (I-like, star-like and so on) is presented in Fig. 6 (I-like). Upon folding of long intronic RNA two processes are substantial: rebuilding of nascent RNA (annealing and re-annealing) and formation of slips. Due to the high AT-composition of introns, short double-stranded (ds) AT-rich fragments can re-anneal at room temperature [44], while longer or GC-rich fragments can be rearranged to a lesser degree. Fragments with dsAlu cluster can form a clip through annealing, which stabilizes the structure (with high thermodynamic preference), as seen in Fig. 3-7 marked in orange. Other possible types of clips are associated with protein binding and long complementary oligonucleotides. The presence of clips, as usually exhibited by dsAlu cluster, mainly determines the existence of reproducible substructures, such as some of Bn branches.

**Fig 3.**
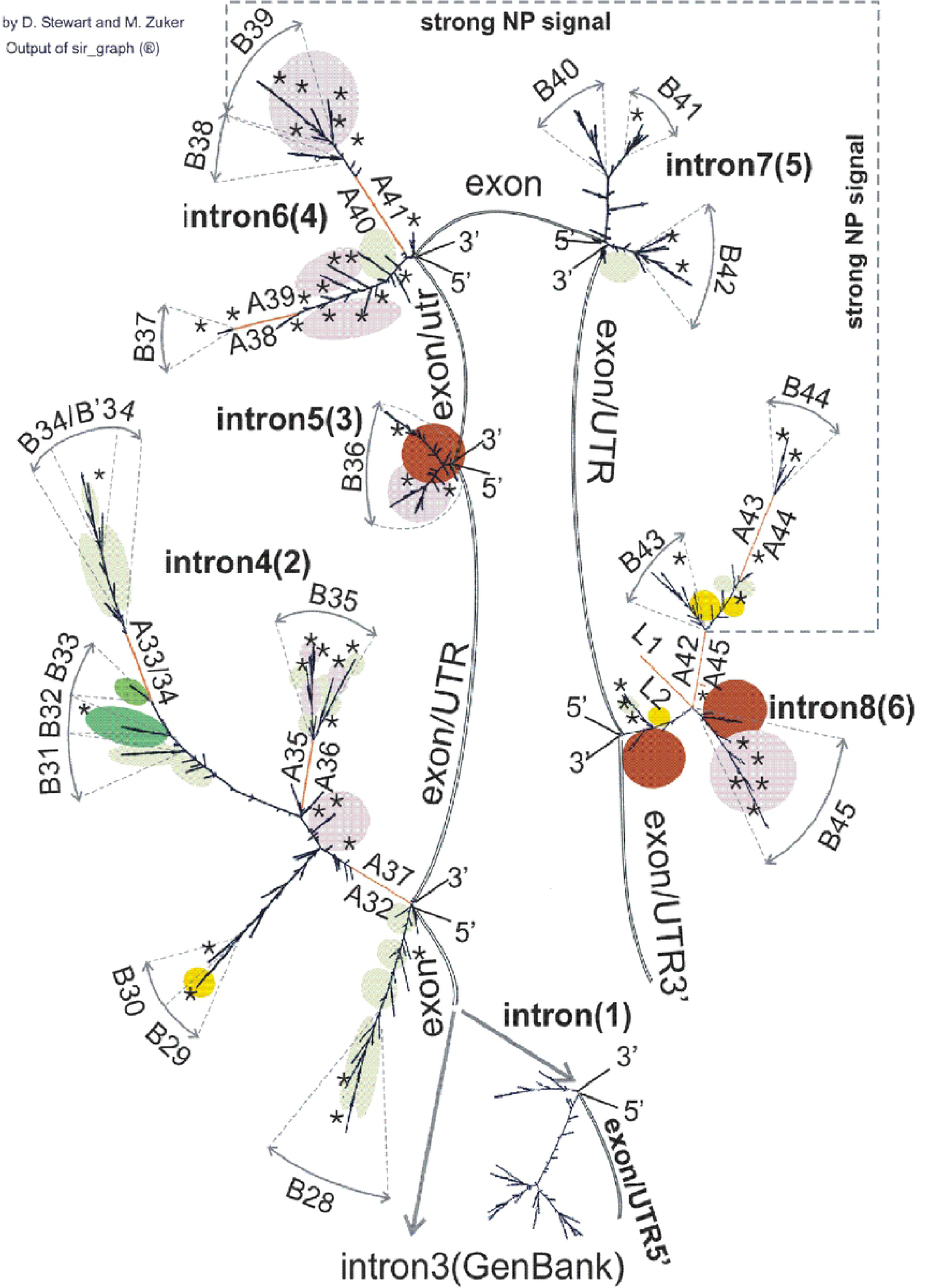
Secondary structure of intron 3 (GenBank) transcripts (intron 3 – intron 9, Gene id) and intron 1,2 (GenBank). It is the basis for mapping of RNA-binding protein sites for the part of variant 1,2 of GABRB3 in the recursive splicing version. Assembly of intron 3 – intron 9 (Gene id) corresponds to intron 3_GenBank.

**Fig 4.**
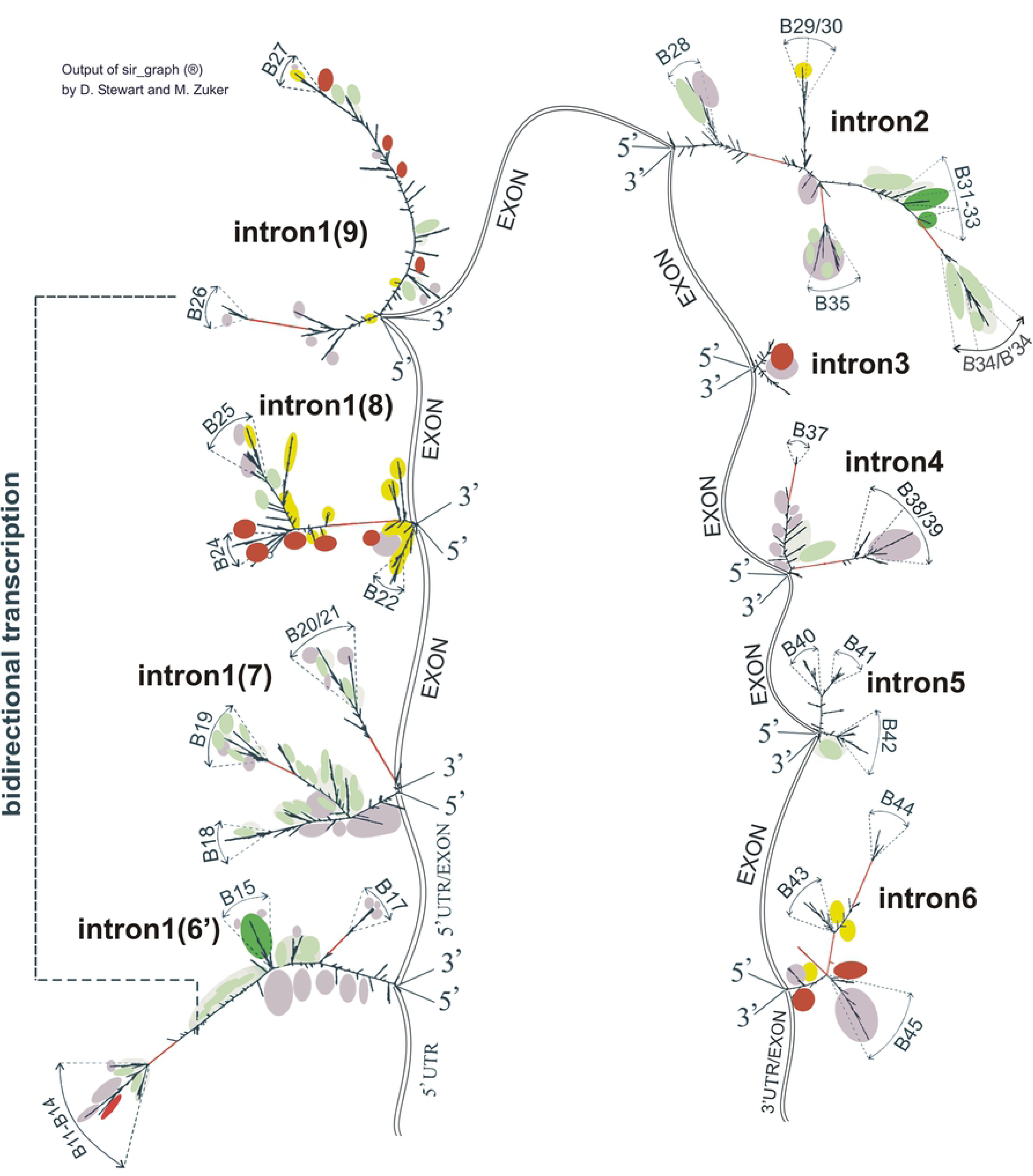
Secondary structure of intron transcripts as the basis for mapping of RNA-binding proteins sites (variant 4 (core-portion)). This variant also constitutes a part of variant 1,2 of GABRB3. Intron enumeration is done according to GenBank for variant 1,2 (enumeration for variant 4 is shown in parentheses). Colour spots are as in Fig. 3.

**Fig 5.**
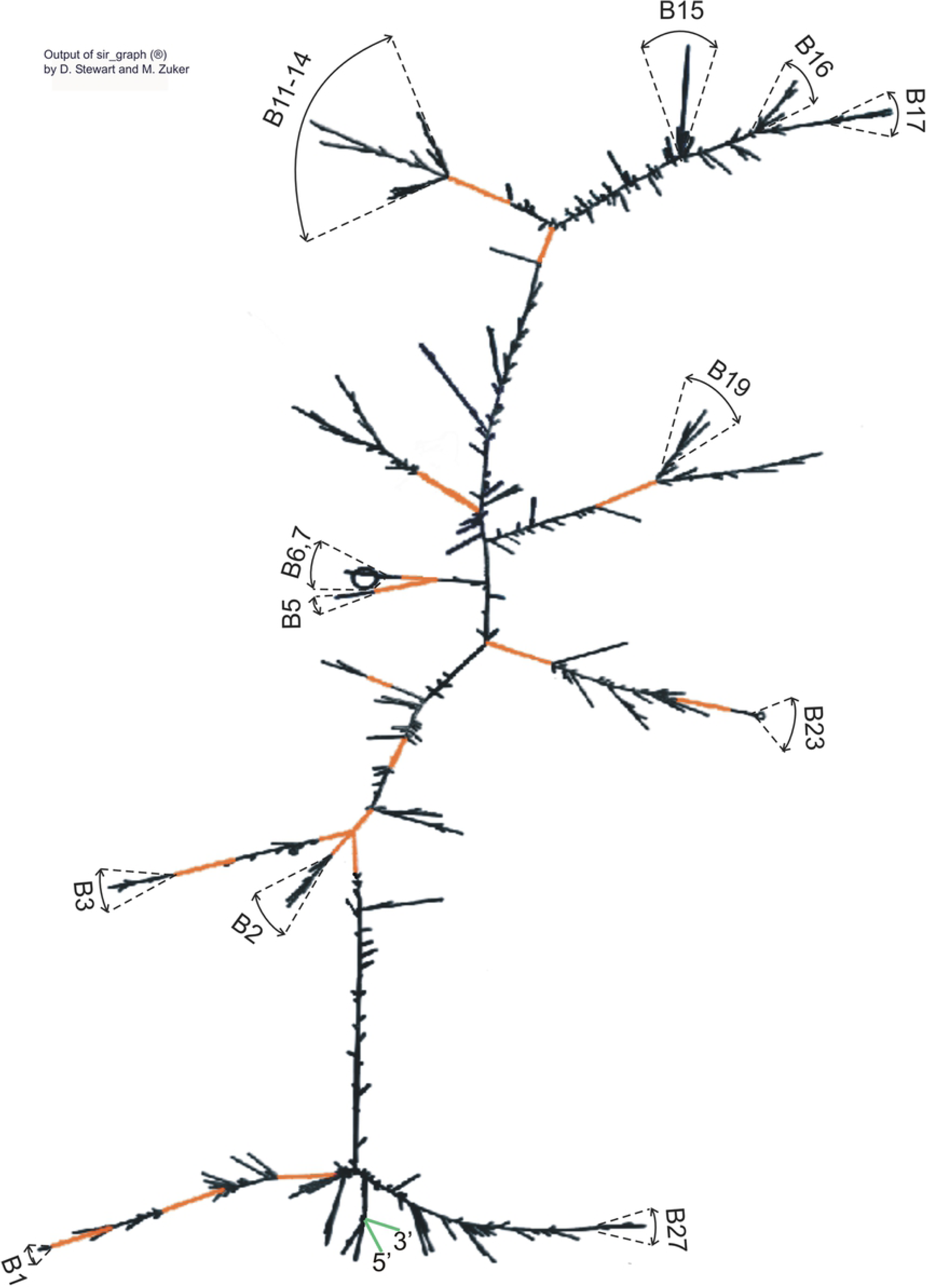
Secondary structure of intron transcripts (GenBank) as the basis for mapping of RNA-binding protein sites (variant 3 of GABRB3). Enumeration of introns corresponds to constitutive splicing (GenBank), in parenthesis, correspondence to Gene id enumeration of long intron subdivision as in Fig. 3, intron 1 (6’) is a truncated variant of intron 6 (Gene id, part of intron 3, GenBank). Colour spots indicate association with RNA-binding protein sites are as in Fig. 3.

**Fig 6.**
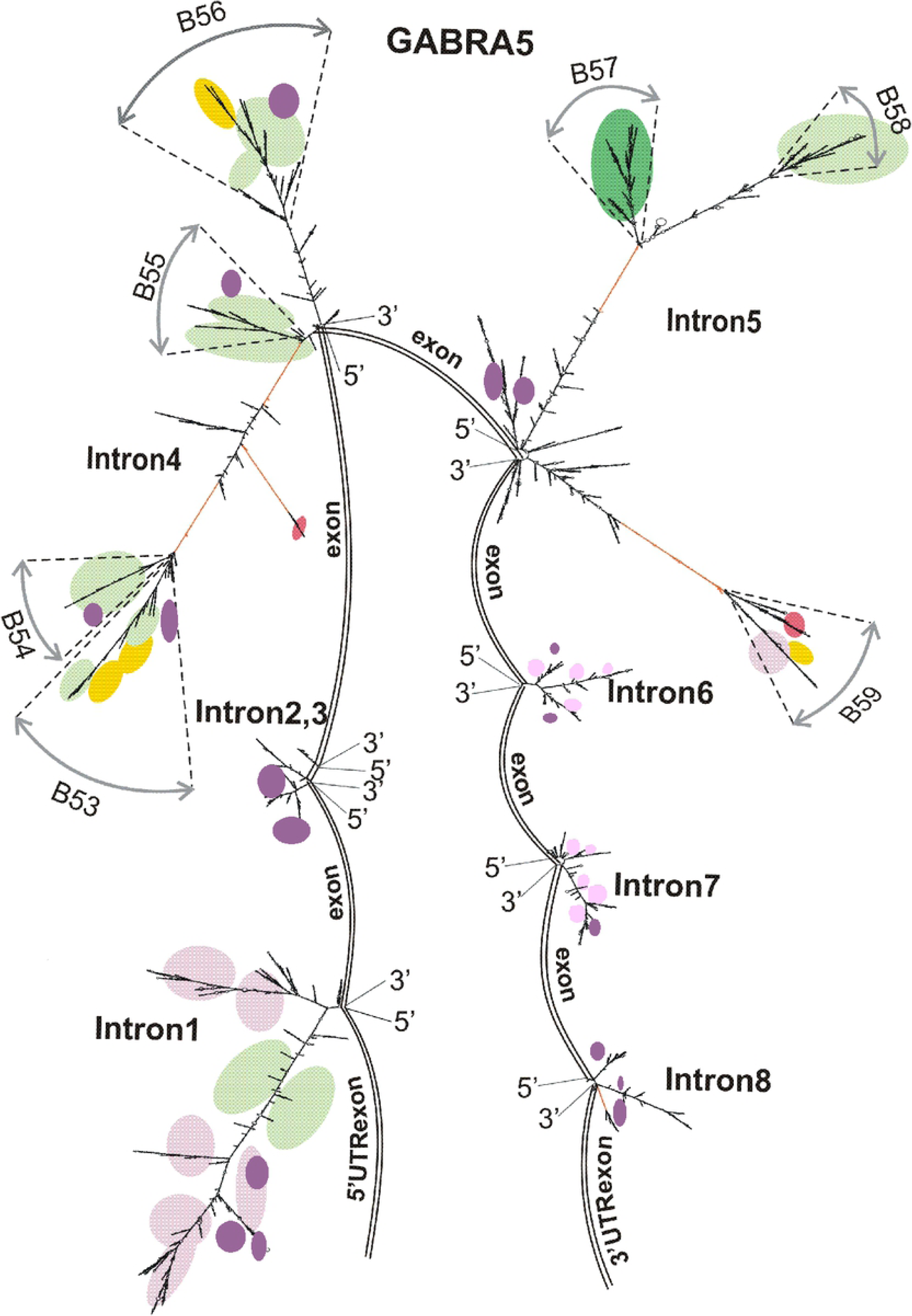
Example of secondary structure of intron transcript (intron 3_GenBank) as a part of variant 1, 2 of *GABRB3* (without recursive splicing). dsAlu is shown in orange, Bn - reproducible branches.

**Fig 7.**
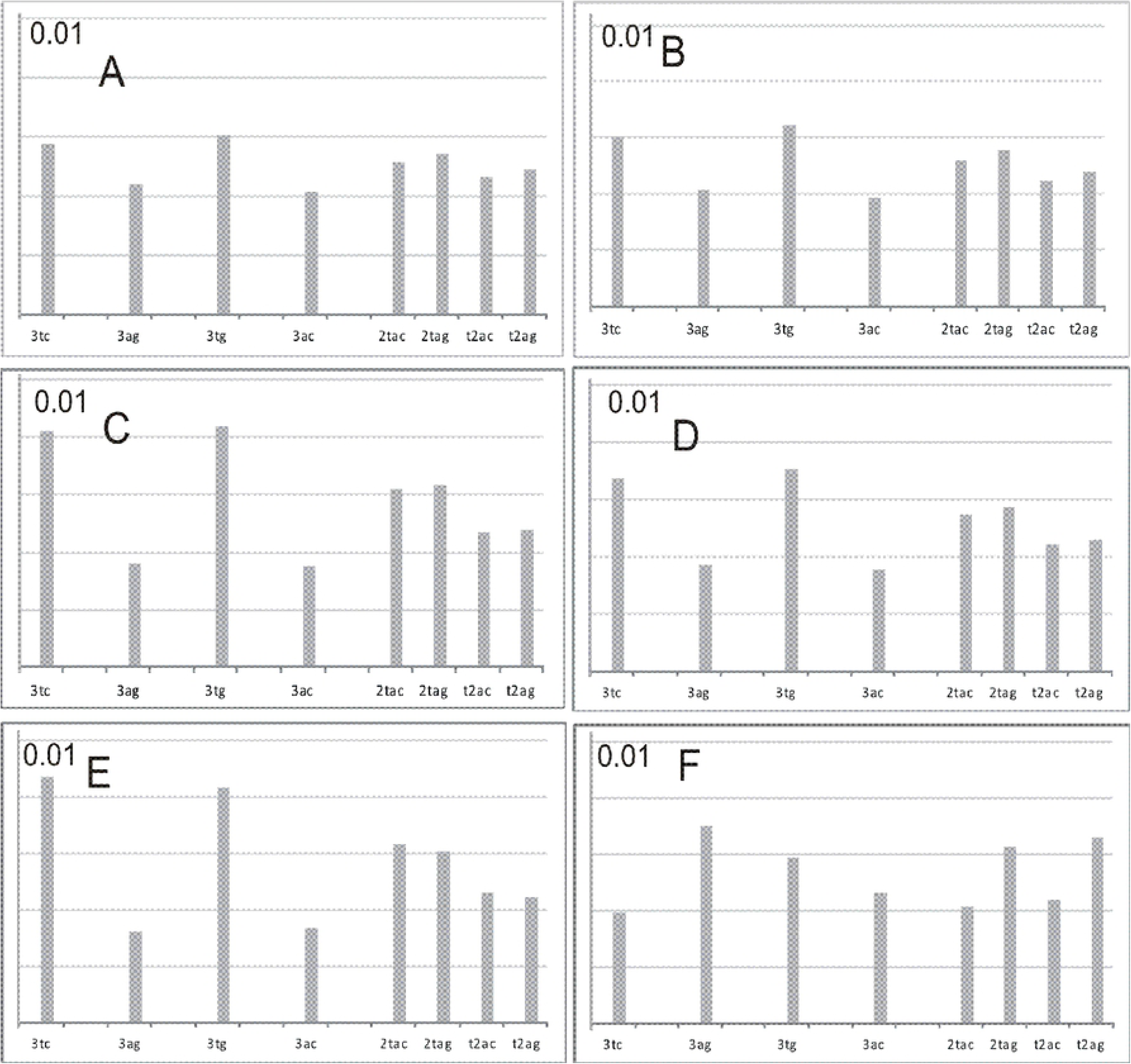
Secondary structures of intron transcripts (GenBank) as the basis for mapping of RNA-binding protein sites (transcriptional variants of *GABRA5*).

## Mapping of protein binding sites

### Serine-arginine protein family

We chose the serine-arginine family proteins that are widely involved in intra-nuclear processes and have been relatively well-studied. *SRSF1* protein (aliases *ASR/SF2, SRp30a, SFRS1*) may represent many functions: (a) an active participant of the spliceosome assembly [55-57]; (b) an exon enhancer-binding protein [58] in the case of double-site purine-consensus (which was not present in the locus) and as a splicing repressor at some locations, particularly in intron sequences [59-60]. The repression may be restricted only to some special situations, and consequently, the analogous sites fail to either activate or repress splicing intron localization in case of intron localization [61]. SR (serine-argenine) proteins, including *SRSF1* and *SRSF2,* are recruited to nascent pre-mRNA, as shown for polytene chromosomes and Balbiany Rings, in the gene-dependent manner and may even relocate during transcription to the more downstream parts of long genes [62,63]. This relocation predisposes them for transport to the cytosol and influences mRNA binding to the ribosome at further stages. Besides, the *SRSF1* protein is a part of NS [64], and this participation may be phosphorylation-dependent, as it regulates alternative splicing [65]; beyond its roles in mRNA splicing, stability, and translation, this protein has other functions related to mRNA-independent processes, such as miRNA processing, protein sumoylation, and the nucleolar stress response [66].

For *SRSF 1* proteins, following the data obtained by the functional UV cross-linking and immunoprecipitation (CLIP) method [49,67] and SELEX [50,51,68], we mapped potential sites for RNA binding on the locus sequence [Fig. 2M-R]. The binding sites derived by the functional method (Fig. 2N) had a consensus sequence SRSASGA (7-mer, S:G or C, R-purine) [49], which was more complicated by its nucleotide diversity than those derived on the basis of hepatitis delta virus genome sequence, which had a purine-rich consensus RGAAGARR (8-mer, R-purine) (Fig 2O) [50] or by the SELEX method (Fig. 2P), which was represented by RGAAGAAC (8-mer, R-purine) [51]. Despite the differences in consensus, the overall numbers of combined binding sites are presented for completeness in Fig. 2Q for the (-) strand and Fig. 2M,R for the (+) strand.

Our mapping data are presented in two forms: mapping of binding sites (a) on the primary sequence of the locus in linear representation, and (b) on the secondary structure image (2D) of intron RNAs. Some branches, namely, B9, B12-14, B39, and B45, are enriched in these sites. A much greater extent of enrichment has been found for the inter-branched spaces: 5’-A1, interA5-A7, inter B17-18, interB26-B27, inter B38-A41, interA35-A38, L2-5’ (Fig. 2M-R, Fig. 3-5,7, as violet spots for site density higher 1.5 motif/knt and as a star * in Fig.4) (*GABRB3*), whereas B42 and B44 are enriched to a lesser extent. For the longest pre-mRNA variant of *GABRB3* (Fig. 1L), significant peaks in the (-) strand are also present in the region intersecting with *GABRA5* gene. In the case of 2*D mapping, the set of violet spots (Fig. 3-5,7) coincides predominantly with intron 6,7 (Gene id, middle portion of intron 3, GenBank) as well as with the 5’end of intron 3 (GenBank) and indicates enrichment with *SRSF 1*-binding sites. It should be noted that exons are not highly enriched in *SRSF1* binding sites; they are at the same average density level as introns. The density of binding sites for a long intron 3 (GenBank) is equal to 1.2 units/knt; for intron 4 (GenBank), it is approximately 0.4 units/knt; and the overall number of binding sites for intron 3 (GenBank, f2 fragment in Fig. S1B, Fig. 2T) is several times higher than the number of binding sites for the core portion of *GABRB3* gene (f1 fragment, Fig. S1B, Fig. 2T). The introns 5, 6, 7, 8 (Gene id, part of intron 3, GenBank) (density ∼1.21 units/knt) are more enriched in *SRSF1*-binding sites than intron 4 (GenBank). Introns 5-8 (GenBank, *GABRB3*, Fig. 2Q, Fig. 4) also have significant levels of *SRSF1* binding sites.

For *GABRA5* gene ((+) strand)) a strong peak I is located upstream of TSS (*in silico* predicted chr15.140.2 by Genscan), other strong peaks II, III are in intron 1, at the boundaries of introns 2,3 and in intron 6-8 (they alternate with peaks in the (-) strand). For intron 1 (GenBank, *GABRA5*) the density is equal to 1.74; for introns 4 and 5 (GenBank, *GABRA5*) it is equal to approximately 0.40 and 0.78 units/knt, respectively; for introns 6-8, it is equal to approximately 1.7 units/knt.

In other words, the beginning and the end portions of *GABRA5* gene, as well as the middle and the 3’ end portions of *GABRB3* gene, are enriched in *SRSF1* recognition sites. Accumulation of recognition sites at the 3’ end portion of both genes may be useful for RNA processing in this portion, which has a high exon density.

Extended first introns with high total amounts of *SRSF1* binding sites, as in the case of *GABRB3* and *GABRA5,* because of the length and accessibility for scanning are more likely to reach the borders of the inter-chromosome GC, incorporating *SRSF1* (important for spliceosome assembly) and *SRSF2* for recruiting them or binding freely dispersed protein molecules, thus raising their local concentration the in gene vicinity, and therefore, first introns are prone to postTS. Intron 3 (GenBank, *GABRB3*) enrichment in *SRSF1* recognition sites can serve as a storage device for an downstream area with densely located introns and exons in the case of postTS. Altogether, this can lead to an efficient processing of pre-mRNA.

According to the proteomics data (GeneBank) [69], the density of *SRSF1* protein in brain is at a medium level (21.9 RPKM) (max level 44.79, min 4.4 RPKM), and the manifestation of function may be stronger in tissues with a higher level.

Another protein, *SRSF2,* from the same –serine-arginine family is believed to be present in NS (granules), involved in alternative splicing, appear in the differentiation of stem cells and play a role in transcription pause release in Drosophila and mouse, and, together with *SRSF1,* in mammals [70,71]. A disease-associated mutation in *SRSF2* gene results in mis-regulated splicing by altering its RNA-binding affinities [72].

For the consensuses AGGAGAU and GRYYCSYR (Y-pyrimidine, R –purine, S:G or C) [47,48], explicit predominant localization of binding sites is not observed in the second and third portion of the longest intron 3 (Fig. 2I,J). The mapping shows uniform binding character with the exception of the first third (near the 5’ end) of intron length. The GC-rich site in the promoter zone (*GABRB3*) has intermittent peaks I-IV of potential binding sites (Fig. 2J, Fig. 3, as dark violet spots for site density higher 4 motif/knt). These peaks are also observed in introns 1, 4 of *GABRA5* for the longest variant of *GABRB3* ((-) strand) (Fig. 2L). Analogous zones of intermittent peaks in introns 1 and 4 of *GABRA5* gene for the (+) strand are represented by peaks I-III (Fig. 2L). In the core portion of *GABRB3* (variant 4) (Fig. 2J) and inter-gene portion between *GABRB3* and *GABRA5* (Fig 2J) the signals of potential binding sites are scarcely present. The peaks of potential SRSF2 potential binding sites are predominantly present in the downstream regions adjacent to 5’-ends of both genes, which may be associated with their role in the transcription pause release.

The gene expression according to Proteomics [69] (hppt://ncbi.nlm.nih.gov/gene) for brain is estimated as 30.1 RPKM, which is about the average level (max level 91.7 and min level 8.8 RPKM).

For *SRSF5* protein (aliases *SRp40, SFRS5*), an important function is related to regulation of a significant factor switch through alternative splicing, and this is associated with a great variation in its concentration in utero during pregnancy [73]. The RNA recognition sites of *SRSF5* protein are presented by ACDGS (D:A,G, or U,S:C,G) [49]. These sites are predominantly localized in the regions that are relatively free from other proteins. On a fairly uniform background (*GABRB3*) as a whole, the lateral zones, namely, intron 8 (Gene id, part of intron 3 (GenBank)) and, to some extent, intron 2 (Gene id, part of intron 3 (GenBank)), as well as intron 8 (GenBank) (second portion of core-gene) are enriched in *SRSF5* protein-binding sites (Fig. 2G, Fig. 3-5, as brown spots for site density higher 10 motif/knt). For *GABRA5,* some strong peaks are present in introns 1,4,5 (GenBank) (Fig. 2H, (+) strand), and in inter-gene region, binding sites are scarcely present.

For an easy description, we highlight fragment f1 as containing short introns, exons and 3’UTR, f2 -long intron, f3 – *GABRB3* and *GABRA5* inter-gene region, f4 - long intron of *GABRB3* gene in accordance with the *in silico* predictions, and f5 – *GABRA5* gene, f3+ f4 + f5’ intron transcription variant of *GABRB3* gene (end-to-end across the *GABRB3* and *GABRA5* genes, active in retina). Quantitatively, the *SRSF 1* signal density in f2 and f3+f4+f5’ for long introns (S1 Fig. 1, S1) exceeds that for f1 (data for the gene-core), especially, when integrating over the entire length of *GABRB3* gene, when it becomes obvious that the main share falls on long introns, as if they collect *SRSF 1 protein molecules*. The density values are approximately at the same level. The same situation is observed for *SRSF 2, 5* proteins. Somewhat elevated levels of *SRSF 1, 5-*binding sites are present in the area adjacent to the 3’-ends of both genes.

### PTB protein binding

Other important poly-pyrimidine binding proteins *PTB P1* and its paralog *PTB P2* (expressed exclusively in brain and especially in neuronal precursor cells) are multi-functional proteins [74]. *PTB P1* functionally mediates the formation of RNA loops [75] and also competes with *U2AF* for binding to Py-tracts (PPT) [76, 77]. *PTB P1* can influence alternative splicing and exon-skipping in certain cases (for example, exon skipping in *GABRB2* gene in non-neuronal splicing extracts) [53, 54, 78], although the *PTB P1* level in the brain, compared to other tissues, is sufficiently low (Proteomics data [69]). Its RRM1 (RNA recognition motif) binds to single-stranded RNA; while RRM1 and RPM2 remain independent in solution, RRM 3 and RPM4 may interact with each other producing a single globular protein moiety [79,80].

The RNA sequences for PTB binding contain 15–25 pyrimidine bases, with a preference for special pyrimidine tracts containing UCUU, UUCU, (C)UCUCU [52,53,81]. The occurrence of the UCUU/UUCU motif is significantly higher than of the (C)UCUCU motif, and as usual, the motif is co-localized with more nonspecific Py-tracts. We mapped these tracts along the 15q11-12 locus. In the long intron 3 (GenBank, *GABRB3*), they are localized in the central part in one-dimensional representation (Fig. 2T) and in 2*D representation (Fig. 3-5,7, green or dark green spots, green spots for tracts density higher 20 motif/knt, dark green ones for density higher 25 motif/knt), namely, in the strong peak B15 and in the weaker peak B11, as well as in inter-branch spaces (interB14-B15, interB15-B16). For the core-part of *GABRB3* gene (Fig. 4), such mapping revealed strong peaks ?32-33, B34/B34’, B35. For a long intron 4, 5 of *GABRA5* gene (GenBank), the peaks in one-dimensional representation (Fig. 2U) correspond to branches B55, B57, B58, also shown in 2*D representation (Fig. 7). For chr15.140.2 intron of *in silico* predicted transcription variant, a strong peak of specific Py-motives up-stream of the 5’-end of *GABRA5* annotated transcripts was also mapped. For further elucidation of the degree of incorporation of specialized PTB-binding Py- motives among nonspecific Py-fragments, we assessed the amount of 15-25 nt Py nonspecific fragments incorporating the specific Py-motives. A large portion of specific PTB-binding Py-motives is dispersed outside of continuous fragments containing 15-25 pyrimidines.

In the B11 branch, there is only one perfect Py fragment (>15 nt) with specific Py-motives, in B15 - 1 Py-tract (>15 nt) with specific Py-motives, in B31-33 – 4 tracts, in B34 - 8 Py tracts of 15-nt fragment, in B35 - 4 of 15 nt, in B55 - 1 of 15 nt, in B57 - 3 of 15 nt, in B58 - 1 tract longer than 15 nt with specific Py-motives (Fig S2).

Green spots dominate the upper part of the picture (Fig. 3), that is, in the centre of the long intron 3 (GenBank). In intron 4 (GenBank), the intensity of Py-rich motives and tracts is higher than in intron 3 (GenBank) and is more concentrated. This increased level refers both to the overall number of motives and to the number of almost full-size fragments (>15 nt) enriched in specific Py- motives. The level outside of this zone is quite low. A middle portion of *GABRA5* gene (introns 4, 5) also contains strong peaks.

Pyrimidine tracts in introns of *GABRB3* and *GABRA5* genes are quite far away from the splicing sites in terms of their localization, so they are unlikely to affect exon skipping, and their role in the remote localization of splicing sites remains unclear. However, their influence on organization of long RNA loops by PTB P complexes with their repressor role in a weak and regulated exon splicing in a tissue and differentiation stage-dependent manner, which needs cofactors, cannot be excluded.

### hnRNP

*hnRNP L* protein binds (CA)n, where n is approximately 30 and is localized at a certain distance from the 3’-splicing sites inside exons, where it serves as a splicing enhancer [82]. In most other cases, it serves as the silencer of splicing or an enhancer depending on its binding site proximity to the alternative 5’-splice site [83]. Not only regular CA repeats are recognized with high affinity, but so are certain CA-rich clusters. In our study, *hnRNP L* mapping was conducted in accordance with SELEX data [83] for regular CA repeats that are marked by yellow spots. Long (CA)n repeats (n>30) are not encountered in the locus. Yellow spots in 2*D secondary structure image (Fig. 3-5, 7) correspond to oligomers, and their localization favours the silencing potential of splicing according to published data (83).

For *hnRNP G* binding sites, there is a preference for CCA repeats [84]. The B6 branch of the long intron 3 (Fig. 3) has sufficiently long repetitions of similar sequence. As *hnRNP G* and *hnRNP L* binding sites may have some overlap, they are both labeled by yellow spots (Fig. 3-5,7, as yellow sports for density higher 22 motif/knt). Note that *hnRNP L,G* proteins are present in the brain at the level below the average, compared to the spectrum of other tissues according to the Proteomics resource [69]. The CCA trinucleotide is a part of CCAT repeat, and both are encountered in B6. CCAT is the binding site of *YY1* transcriptional factor that binds both DNA and, with a lower specificity, RNA as well.

*hnRNP A1* performs many roles. It is known that *hnRNP A1* and *SRSF1* compete with each other for implementation of alternative splicing [86)]. *hnRNP A1* also interacts with telomere sequences [87,88], associates with granules and accelerates annealing of single-stranded substrates [89]. The level of this protein in the tissues is high in accordance with many functions, whereas in the brain, it is below the average level. Mapping of *hnRNP A1* high-affinity binding motif TAGGGA/T [46,85] shows many potential binding sites, dominating in the GC-areas, and outside of the long intron the binding sites are as frequent as in the long intron (Fig. 2C-F).

*hnRNP C* is highly concentrated RNA-binding nuclear protein (in brain, approximately 48.8 RPKM, max 82, min 7.76 RPKM), it recognizes 5U and 4U [45]; however, not all potential binding sites are occupied, and upon mapping 5U tracts (Fig. 2A,B), its plot approximately follows *PTB P* curves with some variations. We also mapped the *hnRNP C* binding site (5U) at the locus in 2*D representation and accomplished a computer simulation of the folding process by replacing 5T with 5N in order to take into account the effect of high nuclear density on the folding result. The comparison results relate to some changes in non-reproducing branches (data not shown). An influence of *hnRNP C* protein on the details of substructures requires separate consideration.

Favouring the requirement of long intron protection from premature cleavage and polyadenylation [17-18], our results present the evidence of an increased density of polyadenylation signals for intron 3 (GenBank, *GABRB3*)(density of polyadenylation signals, 0.95 unit/knt) in comparison with the density for other introns (0.6 – 0.7 unit/knt). The maximum density of 27 is for A8 (Alu) in the polyadenylation site cluster. For *GABRA5* gene introns, the density of polyadenylation signals is in the 0.4-0.5 unit/knt range, and for both genes, the first introns have maximal density level.

In addition, despite the high level of *Drosha* (RNase III) in the brain [69], it should be emphasized that the locus does not contain a *Drosha* pre-mRNA substrate for processing into miRNA. Additionally, this locus does not contain *REST*-binding sites (*REST* is a transcriptional repressor of neuronal genes in non-neuronal tissues).

### Enrichment of simple nucleotide tracts and its potential for branch interaction with ncRNA Malat1

Prevailing dispersed distribution of short (4-6 nt) Py-motives recognized by *PTB* proteins prompted us to investigate oligonucleotides frequencies in this locus. We placed an emphasis on tetramers of simple nucleotide sequences, as they are the shortest simple sequences that are recognized by RNA-binding proteins, and they have an optimal length for complementation of single-stranded loops of secondary structure (kissing). More complicated by sequence composition and accordingly less-frequent tetramers encountered in AT-rich introns are too complex for simple analysis.

Proteins interacting with Py-tracts of RNA are presented by genes *PTBP1, PTB P2, U2AF65* (*U2AF2*). Proteins having the shortest recognition motives are represented by *PTBP1, PTBP2*. They require the UCUU/UUCU recognition motives (*PTBP1, PTBP2* binding sites), and elements of Py-tracts near branch point are recognized by *U2AF65*. To achieve the comprehension of the availability of different nucleotide tracts and their distribution within introns of *GABRB3, GABRA5* genes, we present a preliminary assessment of tetramer occurrence based on nucleotide composition. We accomplished a rough frequency estimate without considering the Markov chain character of nucleotide sequences. According to Fig. 8, introns 5, 6 (Gene id), as parts of large intron 3 (GenBank, *GABRB3*), and intron 4 (GenBank, *GABRB3*) are mostly enriched in 3TC-tracts and 3TG-tracts, whereas introns 3, 4, 7, 9 (Gene id, parts of intron 3, GenBank)) are enriched in 3AC tracts (is not shown). More complicated 2TAC, 2ATC, 2TAG, 2ATG, AT2G and 2G2C tetramers are less representative in these introns, and we did not consider these tetramers with intron 3, 4, 7, 8, 9 (Gene id, parts of intron 3 (GenBank)), with their uniform representation of simple 3TC, 3AG, 3TG, 3TC tracts and more complicated 2TAC, 2ATC, 2TAG, 2ATG tetramers.

Naturally, it is important to search for genomic elements containing Pu-tracts complementary to Py-tracts. These may be the local elements of the same introns as well as distant ones. The remote sequence of ncRNA Malat1 should also be included. According to microscopic studies mentioned earlier [20,27,90,91], the NS (granules) containing serine-arginine proteins and Malat1 are recruited to the peri-chromatin fibril regions (PF), and Malat1 interacts with the nascent RNA [92]. Furthermore, pre-mRNA with Py-tracts can recruit NS for implementation of splicing, and the recruitment and splicing process are realized only in the presence of Py-tracts [91]. Thus, we include ncRNA Malat1 as a component of granules in our oligonucleotide density consideration.

Within the abovementioned introns and according to the mapping results for 15q11-12 locus, we picked out for analysis the apical branches B9, B10, B11, inter B14-B15, B15, B16, B28, B31-33, B34, B35 (*GABRB3*) and B52-B59 elements (*GABRA5*). In Fig. 9A,B, the branches B11,B15,B31-33,B34,B35 and inter-branch fragments B14-B15, B15-B16 are the main carriers of Py tracts according to the density and overall number representation. In Fig. 9A,B, for branches, inter-branch fragments and branch sub-structures the overall count of tracts for the full-length and density count per 1 knt of item length is equal to count(Py) = number(TTCT) + number(TCTT) - number(TTCTT), count(Pu) = number(AAGA) + number(AGAA) - number(AAGAA).

In Fig. 9A, the differences between count(Py)-count(Pu) for overall count of tract per fragment length are presented, as well as differences in their density per fragments length (Fig. 9B). As follows from difference plot between Py and Pu, the branches B10, B14-B15, B15 (intron 3, GenBank, *GABRB3*) and especially B28, B31-B33, B34’, B35 (intron 4, GenBank, *GABRB3*) are enriched in dominating Py-tracts over Pu- tracts, the same applies to B52, B54-B58 (*GABRA5*), and on the contrary, Malat1 is enriched with Pu-tracts versus Py-tracts, especially the Malat1-2 fragment. Analogous plots are depicted for pairs of CTTT/TTTC versus AAAG/GAAA tracts, as well as TTGT/TGTT versus AACA/ACAA and GTTT/TTTG versus AAAC/CAAA. Analysis for the Py and Pu pairs (CTTT/TTTC and AAAG/GAAA) confirms the domination type deduced for the abovementioned Py (TCTT/TTCT) and Pu (AGAA/ AAGA). For TTGT/TGTT and AACA/ACAA tetramers, the difference plot (Fig. 9C) reveals the domination of TTGT/TGTT over their counterpart AACA/ACAA throughout most of the sequence collection (e.g., B9, B10 and so on, including Malat1), and this observation correlates with an active participation of UG-rich and UA-rich motives in double-stranded stretches of the whole RNA folding structure, and in addition, it is an important for another discussion.

In more detail, for intron 6 (Gene id, part of intron 3, GenBank) the density of Py-tracts (TTCT/TCTT) is 13.4 units/knt versus 11 for Pu (AAGA/AGAA) tracts, for intron 4 (GenBank), the density of Py-tracts is 15.7 unit/knt versus 8.6 for Pu tracts. For example, for B15, the density of Py-tracts is almost twice as high as for Pu-tracts, e.g., ρ(Py) ∼ 15 units/knt, ρ(Pu) ∼ 8 units/knt, (the highest concentration of Py in intron 3, GenBank), for B31-33, ρ(Py)∼29, ρ(Pu)∼5, B34 ρ(Py)∼20, ρ(Pu)∼10, B55 ρ(Py)∼20, ρ(Pu)∼5, B57 ρ(Py)∼30, ρ(Pu)∼4, B58 ρ(Py)∼23, ρ(Pu)∼5 unit/knt. The density in intron 4 (GenBank) is greater than that in intron 6 (Gene id) and in the entire intron 3 (GenBank). Also we determined that UUCU/ UCUU tetramers in B10, B11, inter B12-B15, B15, inter B15-B16, B16 are present at 50% in single strand (ss) state (in detail, for B15 near 35% are in ss state and ds state, and 30% are at the junction of ss and ds states); for B’34, approximately 43% are in ss state; for B55, about 45% of motives are in ss state; for B57, about 58%; and for B58, approximately 56% are in ss state. These findings indicate that the ss state density for Py-tracts in intron 4 is greater than in intron 3 (GABRB3), and the highest Py-concentration in ss state is inherent to B57, B58 (*GABRA5*). An opposite situation is observed for Malat1, where Pu-tracts dominate Py-tracts by more than 2-fols. An average density of Pu-tracts reaches 17 unit/knt, of Py-tracts, 8 units/knt for Malat1, and particularly in the middle part of Malat1-2, the maximum density reaches 31 unit knt. In Malat1-2 fragment, AGAA/AAGA tetramers are present at 45% in a ss-state.

In the local interaction between ss of branch loops of Py and Pu types especially for interB14-B15, B15, B31-B33, B’34, B55, B57, B58, the equilibrium is significantly shifted to the prevalence of Py tracts, and local interaction between the branches/inter-branches is unlikely to lead to full compensation of the redundancy of Py-tracts, and conversely, for Malat1, the equilibrium is largely shifted to the prevalence of Pu- tracts. The coincidence of Pu-tetramer concentrations in Malat1 fragments and Py-tetramer concentrations in *GABRB3* and *GABRA5* highlighted the most significant fragments as well as ss-state portions. First, this means the ability to interact in complementary fashion at the nucleotide level through the mating loops, namely, the formation of tertiary structure elements between coding RNA and Malat1. For *PTB* proteins, the structural specificity of pyrimidine tracts was confirmed [93], suggesting that the preference refers to unstructured, ss strand variants in main motives. Tracing the ss loops with Pu motives of Malat1-2 often resembles a picture of the spatial repetition of ss loops for Py-rich branches of pre-mRNA.

Second, the branches interaction of coding RNA Py-tracts with Malat1 Pu-tracts may be realized indirectly by the participation of proteins *PTB P1,2* and/or *U2AF65,* and the influence of *hnRNP C* also cannot be excluded as well. The protein influence requires further investigation. This interaction may be realized in a tissue-dependent manner, as the concentrations of proteins vary significantly in tissues [69] (data of proteomics for each protein (http://ncbi.nlm.nih.gov/gene)).

**Fig 8.**
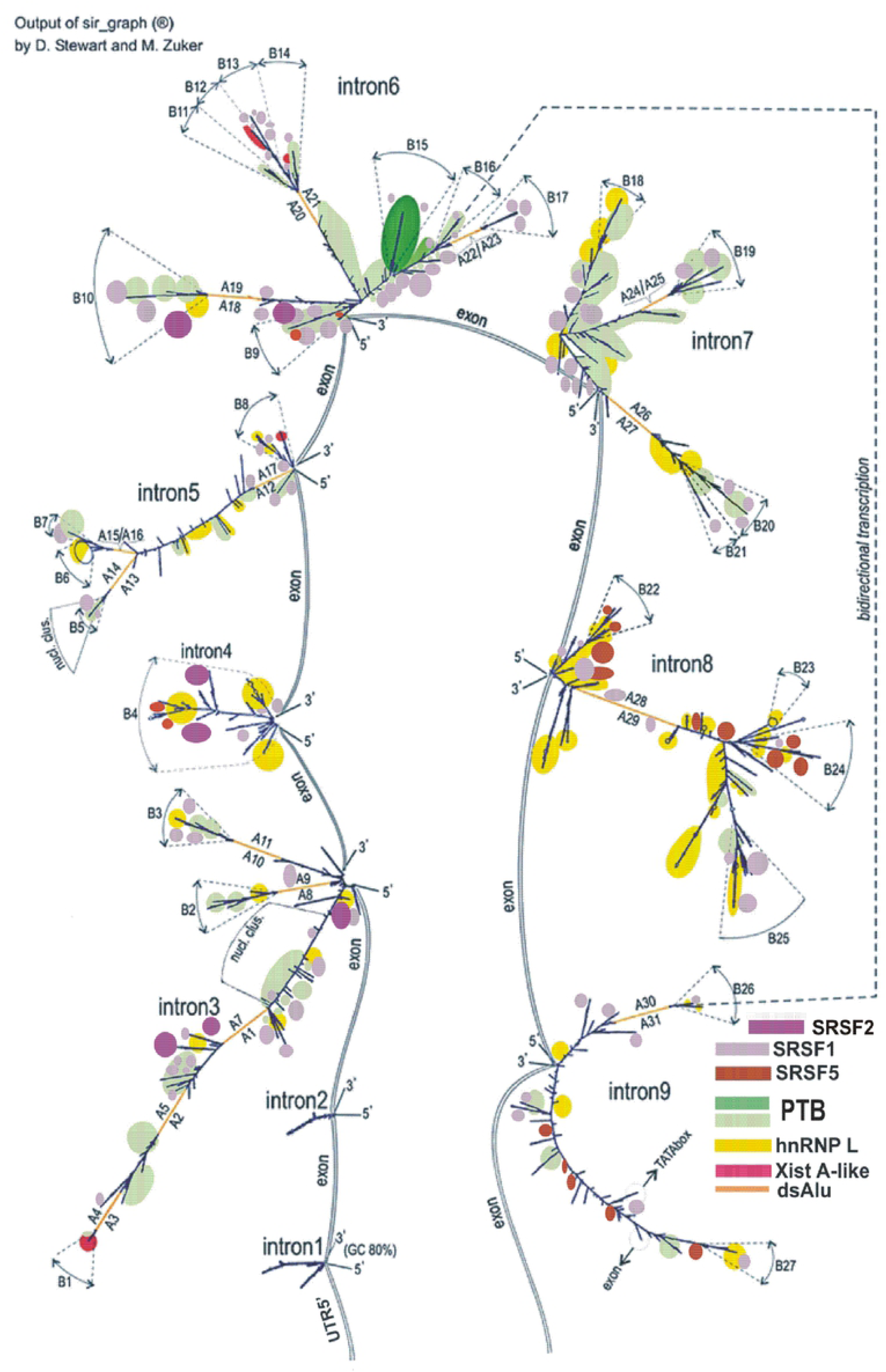
Occurrence of oligonucleotides in introns and Malat1 ncRNA. Intron 5 (Gene id, part of intron 3, GenBank, *GABRB3*). (B) Intron 6 (Gene id, part of intron 3, GenBank, *GABRB3*). (C) Intron 4 (GenBank, *GABRB3*). (D) intron 4 (GenBank, *GABRA5*). © intron 5 (GenBank, *GABRA5*). (F) ncRNA Malat1 ncRNA (EF177381).

**Fig 9.**
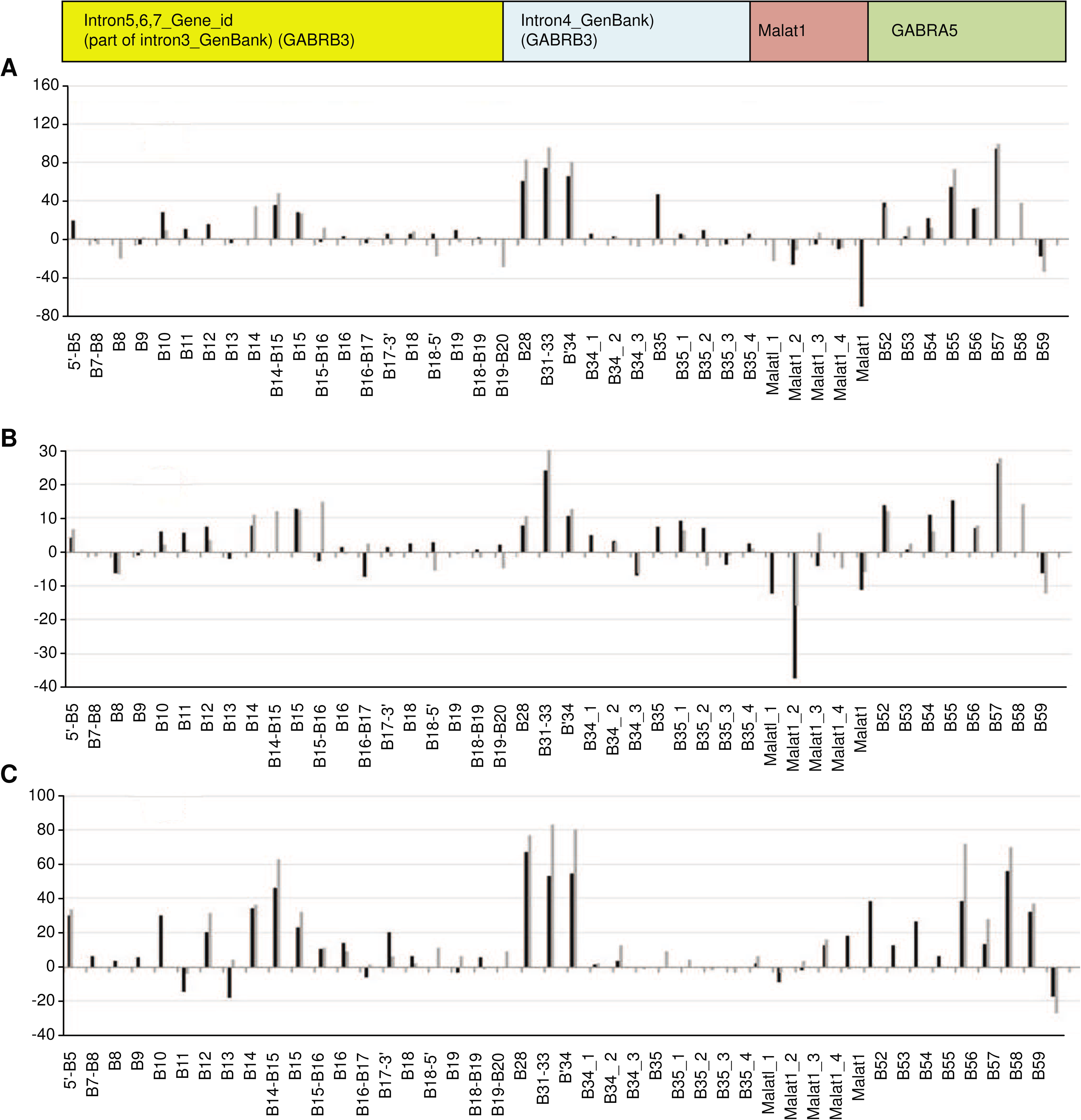
Distribution of the difference between overall number or density for Py and Pu tetramers for some fragments of *GABRB3* and *GABRA5* gene introns. (**A**) (Py (3tc)-Pu (3ag)) difference (overall number of tetramers). (B) (Py (3tc)-Pu (3ag)) difference in density (overall number of tetramers per element length in knt). (A)-(B) TTCT/TCTT and AAGA/AGAA pair is in black, TTTC/CTTT and AAAG/GAAA pair is in gray. (C) (Py (3tg)-Pu (3ac)) difference in density. TTGT/TGTT and AACA/ACAA pair is in black, TTTG/GTTT and AAAC/CAAA pair is in gray.

### Cluster of Alu repeats’

To study the role and structural property of Alu cluster at the beginning of *GABRB3* gene at the level of RNA, we carried out modelling of intron RNA folding in steps. Starting from the minimum (800 nt), the length of fragments was incremented discretely, adding new Alu repeat with an adjacent sequence at each step. This model is equivalent to transcription with long pauses between steps. In a strict sense, this model is different from the native kinetic folding. As shown in Fig. 10, there are 2 chains. The upper one (Fig. 10A-E) extends from Alu1 up to Alu 7. With the addition of Alu 7, further development of the folding following transcription elongation occurs along the lower chain (Fig. 10G-J). In the histogram (Fig. 10M) for the upper chain, the nuclei of dsAlu annealing are Alu 2 (+) and Alu3 (-), and according to the histogram, the interval between them is approximately 1700 nt, whereas the interval between Alu 3 (-) and Alu4 (+) is approximately 400 nt. In the last case, the statistical frequency of annealing is about 2 times higher than between Alu 2 (+) and Alu3 (-). This difference logically follows from the assessment of dsAlu editing rate by enzyme ADAR [94]. It is assumed that this is the case for an average elongation rate for long introns (∼ 3 knt/min). Thus, it follows that the nucleation of annealing for the whole cluster will be generated by the closest *sense* and *antisense* Alu, that is, by Alu 3(-) and Alu4 (+). Most likely, the lower chain is realized for an average rate of elongation inherent to long introns. In the special case of slow elongation or specific pauses, if any exist, the upper chain may by more preferential. After an addition of Alu7 and so on, the lower chain becomes thermodynamically more preferable then the upper chain.

Annealing of the more distant portions of whole chain (Alu5-11) will occur in accordance with the polarity of appearing in the nascent RNA and existing Alu repeats. The lower chain consists of 3 ds-Alu and intermediate fragments containing side branches, having the capacity of stiffening ribs. These fragments (inter Alu1-Alu2, inter Alu4-Alu5, inter Alu7-Alu8, 5’end - Alu1) are enriched in alternating Py and Pu-tracts without a significant type domination. This phenomenon prevents the fragment from folding into a tangle, which would occur for a smooth double-strand nucleotide fragment. After the addition of Alu7 and until the appearance of Alu8, the lengthening of the whole structure occurs at the expense of non-Alu sequences. The subsequent addition of Alu 8-Alu11 does not affect the length of the main telescopic structure.

An important apical branch B1 (∼400 nt) between Alu 3 (-) and Alu4 (+) after annealing has the form close to the structure of A -Xist fragment, and in addition, has the same short oligomers (‘GGAUA’ motif) at the stem-loop junction (for a comparison, A-Xist fragment has fairly evenly distributed 8 repeats [95]), a mutation of this motif induces a decline in binding with Polycomb Repressive complex 2 (PRC2) [96]. This A-Xist directly interacts with the PRC2, which leads to X chromosome inactivation with participation of additional factors [97]. The consensus search for PRC2 binding was unsuccessful and led researchers to suggest promiscuity of PRC2 in association with RNA [98]; however, some preferences were determined (T>A, G>C). For B1 sequence, such preferences are fulfilled (Fig.3, as red spots for ‘GGAUA’ repeat density higher 5 motif/knt). For structural preferences, small RNAs interacting with PRC2 possess 2 stem-loop structures similar to those present within A-Xist RNA, and they have the potential to interact with PRC2, as experimentally established [99]. It is known that PRC2 interacts also with ncRNA and intronic RNAs and, in this regard, our apical structure B1 also has many similarities with 2 stemloop structures as well as with A-Xist structure (compare structures (L) and (K) in Fig. 10LK). These findings are consistent with ideas about the properties of RNA binding site of the PRC2 complex.

The preference of the whole structure due to its length leads to its ability to be exhibited far into the nuclear space, and undoubtedly, due to many degrees of freedom, facilitates the ability to scan the space and cross the area of nucleus as well as to reach distant portion of the same chromosome.

Later in the text, we will show that clusters of significant nucleosome positioning are localized in the downstream area, and this will make the functioning of the complex more efficient in transcriptional silencing in tissues with high levels of PRC2 components. In summary, we can say that in many tissues, the Alu cluster in variant 1,2 (especially in foetal form) may be responsible for one of the possible mechanisms of transcription silencing due to the Alu cluster structure and nearest NP clusters in the long intron. The components of PRC2 complex are not enriched in the brain compared with many other tissues according to the Proteomics data [69], and this associates with expression in brain mostly at a foetal development stage. For a transcriptional variant 3 (truncated variant 1,2 incorporating the deletion of Alu cluster) the expression is allowed in some other tissues, in addition to brain, as mentioned above. However, variant 3 (Fig. 5) also contains some cryptic structural variant of already considered Alu cluster structure, e.g., a prolonged structure with an apical B11-14 substructure (Intron 1(6’)) and special ‘GGAUA’ motives in dense localization with TG-rich content in B12.

**Fig 10.**
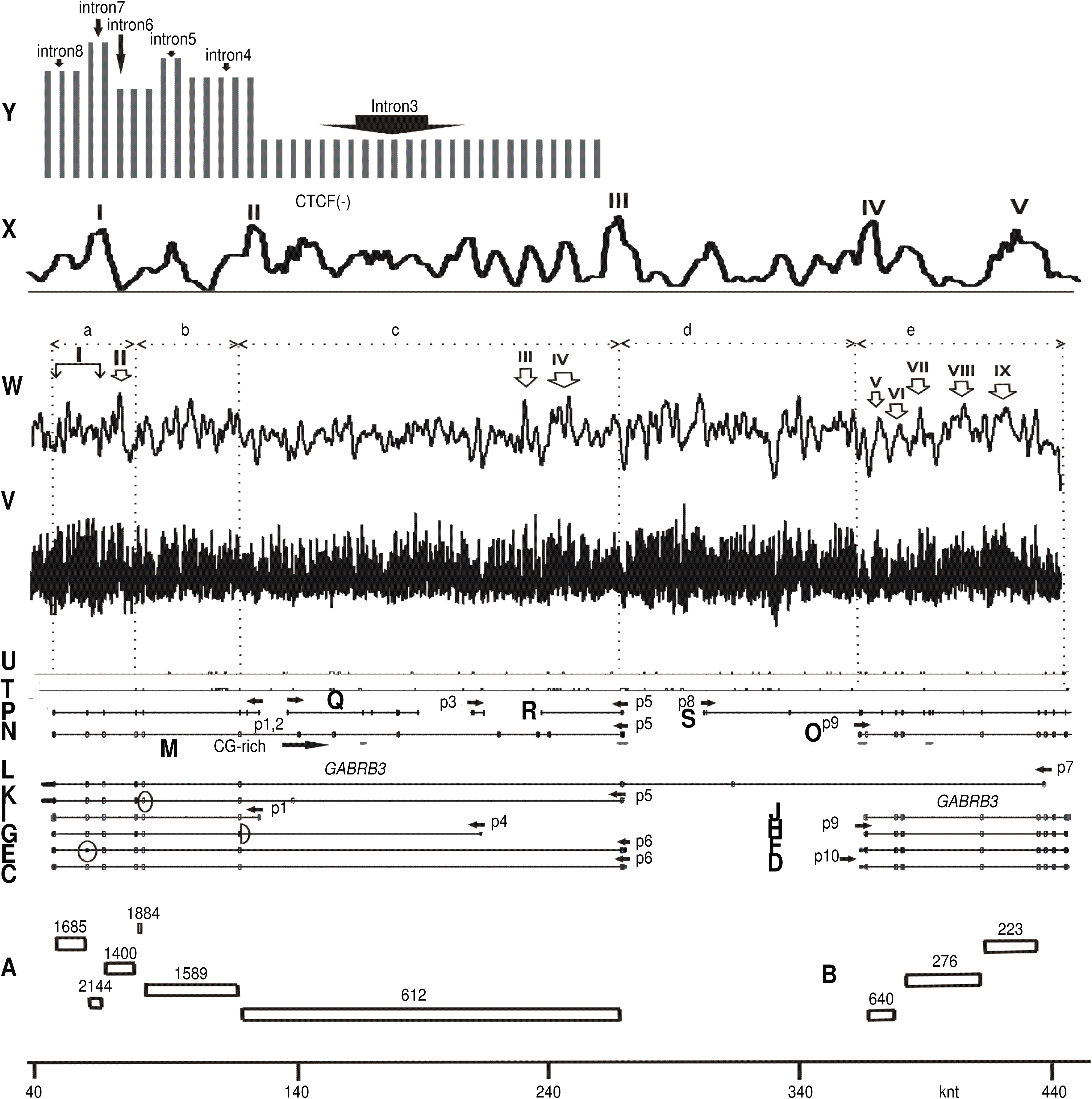
Alu cluster folding at different steps corresponding to the addition of the next Alu repeat with the adjacent sequence. (A) - (F) Upper chain. (G) - (J) Lower chain. (J) Folding of intron 3, Gene id, as part of intron 3, GenBank. (K) Secondary structure of RNA product for branch B1. (L) Secondary structure of A- Xist fragment from [95]. (M) Histogram of the distances between Alu repeats.

### Nucleosome position mapping

In Fig. 11V,W, we suggest two variants of averaging of nucleosome positioning signal (NP) within sliding windows of two sizes (small and large). It is shown that the distinction exists between (a), (d) fragments and remaining portion of the gene (Fig. 11V). Fragment (a) is characterized by alternation of the highest and lowest NP signals, which means the presence of tightly DNA-associated histone octamers isolated by a zero signal from each other, which would probably hamper the formation of a nucleosome cluster and complicate transcriptional elongation. To overcome this hindrance, the participation of the remodelling proteins will be required. These proteins, according to different models, can shift histone octamers by a ratchet mechanism or may lead to the histone octamer eviction [100] and thus eliminate the elongation hindrance. These proteins are part of the SWI/SNF(ISWI) and SAGA complexes [101-104]. According to the Proteomics data [69], these proteins (at least the motor *BRG1* and *SNF2h* ones) have sufficiently high levels in the brain compared with the average level. Fragment (d) corresponding to *GABRB*3- *GABRA5* intergene region also has many peaks and dips in the NP signal (Fig. 11V).

In another situation of II-IX sites the nucleosome clusters may form to be detected when averaging in a sliding window of a larger size. These sites may be transformed to the silencing with the introduction of epigenetic marks that may be generated in the presence of PRC2 complex. It is important to note that, as shown earlier, sites III, IV are localized downstream of a bi-directional Alu cluster together with the apical B1 branch that has some A-Xist–like properties. The oblong secondary RNA structure of the bi-directional Alu cluster with the A-Xist–like spacer may serve as a substrate for the PRC2 complex. This may help the sites III,IV to be transformed to silence state. An additional important observation is that peaks III, IV together with *CTCF* [28] peak III (Fig. 11X) alternate with peaks of *SRSF2* binding sites (Fig. 2J, Fig. 3, dark violet spots) in the region adjacent to the 5’-end of intron 3 (GenBank, *GABRB3* gene). The same situation is identified for *CTCF* peaks (Fig. 11X) and NP peaks V-VII (Fig. 11W) at the beginning of *GABRA5*. It should be noted that *SRSF2* protein helps to release RP-II from transcription pauses [70,71], while strong nucleosome positioning peaks as well as *CTCF* peaks may be the reason of elongation pauses.

In Fig. 11A, Y, the numbers of reads for each intron are presented from the GenBank data. The introns closer to the 3’-end have a higher level of reads than in the middle and more than in the large intron at the 5’-end. This finding is in accordance with the notion that nucleosomes concentrate at the exon edge compared with intron bodies [105]. It is difficult to present unambiguous explanation for this observation. This observation may be related to transcriptional and processing retardation closer to the 3’-end (usually with high exon density), or different levels of transcriptional initiation from the p1, p5 and p6 promoters. This explanation is more likely in the case of the On-line detection system. Another explanation is the possibility of exon-circle formation with the inclusion of introns. The existence of circles formed by exon RNA was detected for *GABRB3* gene [106].

The data in this and previous sections show that the two-level regulation of transcription depends in each tissue upon availability of remodelling or repressive proteins associated with the SWI/SNF(ISWI), SAGA or PRC2 complexes.

**Fig 11.**
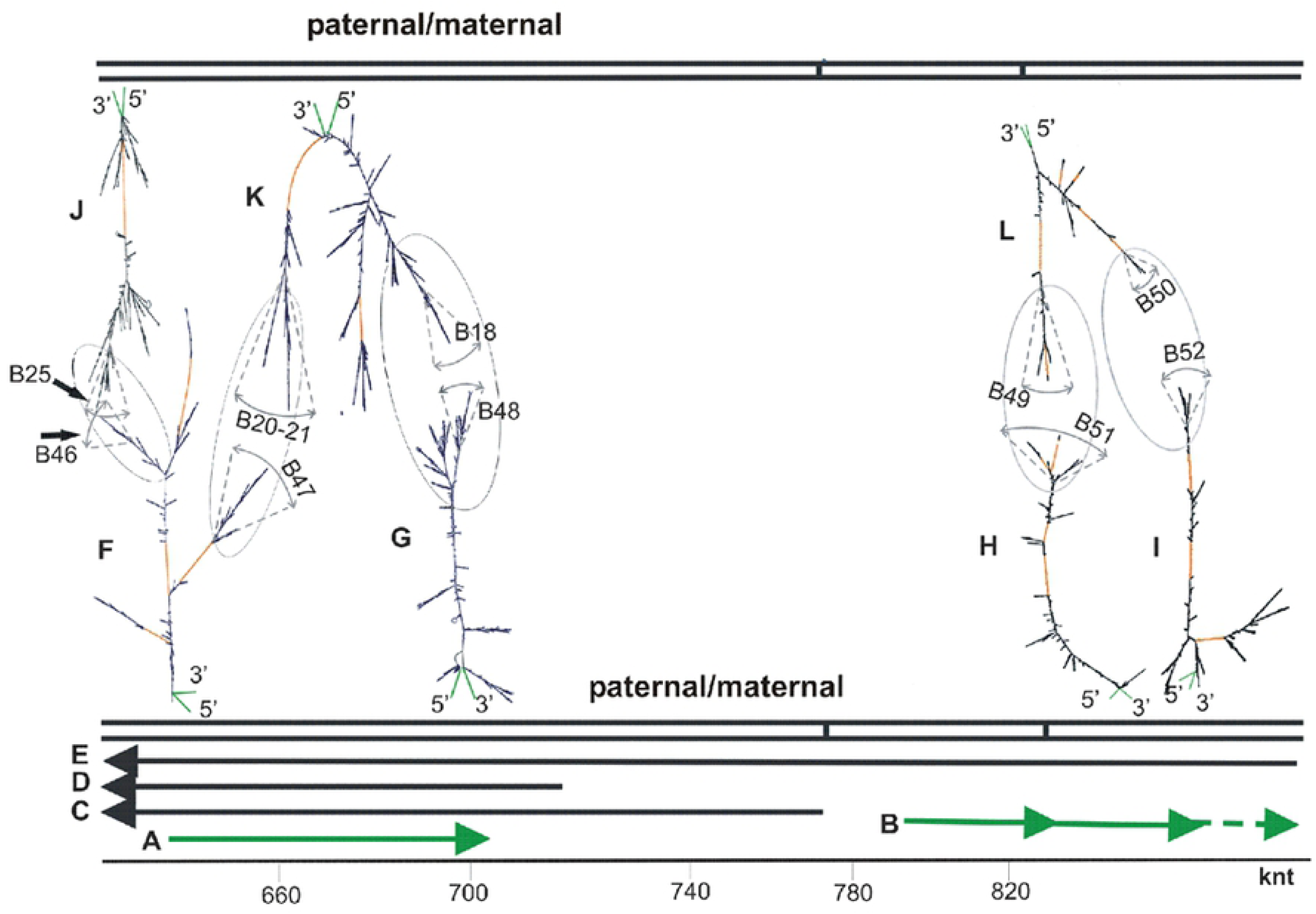
Scheme of the locus, intron reads and mapping of NP and *CTCF* on DNA sequence. (A) Boxplot, number of reads per introns of *GABRB3* gene. (B) Number of reads per introns of *GABRA5* gene. (C) - (U) Scheme of the locus as in Fig. 1. The skipping exons are presented for some transcripts in oval (C),(L), and UTR in half-oval (G). (V) Mapping of NP (averaging in short window). (W) Mapping of NP (averaging in larger window). (X) Mapping of *CTCF-* binding sites. (Y) Histogram of the number of reads per intron length (*GABRB3* gene) https://www.ncbi.nlm.nih.gov/gene/2562.

### Homologous chromosome pairing

In Diptera, the homologous chromosome somatic pairing is widespread and is transcription-dependent [29,107,108]. At least the first stage of this process is in a good agreement with the availability and abundance of bidirectional transcripts in the locus. In addition to Diptera, the homologues pairing was also observed in *Homo sapiens* at some loci, namely, in 15q11-12 [29,109]. Bidirectional transcripts in this locus were shown by GeneBank resources as annotated mRNA and by *in silico*-predicted variants, as well as by availability of bidirectional EST (Fig. 1T,U). We studied a secondary structure of large intron RNA as part of long bidirectional transcripts by UNAFOLD (Fig. 12). Similarly, to the previous cases, the branches may be considered as multiple stable stem-loops substructures with spatially oblong traits. They have approximately the same coordinates on nucleotide sequence (Table S1) and many incidences of complementarity between ss loops in stem-loop structures of branches, namely, complementary sequences of loops in pair of branches, e.g. B25 (intron 8, Gene id, (-) strand) and B46 (chr15.137, intron 1, Genscan, (+) strand), B20-21 (intron 7, Gene id, (-) strand) and B47 (chr15.37, intron1, Genscan, (+) strand), B18 (intron 7, Gene id, (-) strand) and B48 (chr15.137, intron 4, Genescan, (+) strand), B51 (chr15.140, intron 1, Genscan (+) strand) and B49 (part of transcriptional variant CL749803, Fig 1L, (-) strand) as well as B52 (chr15.140, intron 2, Genscan,(+) strand) and B50 (part of transcription variant CL749803, Fig. 1L, (-) strand). Transcription variant CL749809 was revealed at least in retina. In other tissues, this portion of bidirectional predictions of pairing related to CL749809 remains questionable, although the bidirectionality of EST (Fig. 1T,U) supports homologues pairing in wide region. For further justification, we successfully attempted to elucidate the elements of tertiary structure by simulation of ss loop complementary sequences annealing by short oligonucleotides that also confirms the possibility of homologous pairing. As shown, the deletion variants that failed to provide homologous pairing are connected with multiple forms of diseases.

**Fig 12. Scheme of bidirectional transcripts for 15q11-12 locus and images of folding structures of intron RNA.** (A) chr15.137, Genscan, in green. (B) chr15.140, Genscan, in green. (C) *GABRB3* transcript, var1,2, GenBank. (D) Transcript *GABRB3*, var3, GenBank. (E) CR749803 (Table S1). (F) chr15.137, intron 1, Genscan. (G) chr15.137, intron 4, Genscan. (H) chr15.140, intron 1, Genscan. (I) chr15.140, intron 2, Genscan. (J) intron 8, Gene id, part of intron 3, GenBank). (K) intron 7, Gene id, part of intron 3, GenBank). (L) part of intron1, Table S1, transcription variant CL749803, Fig. 1L).

## Conclusions

While studying the thermodynamically equilibrated secondary structures of long intron RNAs, some reproducible substructures are identified. These substructures are the most important results, and they are reproduced in optimal and sub-optimal variants of folding. Many of them are framed by dsAlu repeats and are associated with areas enriched in sites of RNA-binding proteins. For an area in the long first intron adjacent to pre-mRNA 5-end (*GABRB3,* variant1,2) with the cluster of bidirectional Alu repeats, the elongated secondary substructure incorporating a chain of dsAlu repeats (3 units) is identified with an apical stem-loop A-Xist–like branch (A-Xist fragment interacting with PRC2 and pertaining to X chromosome inactivation). The formation of this chain of 3 dsAlu repeats has the preference of occurring at a high elongation rate. While mapping the NP signal on DNA, we also found nearby nucleosome clusters on the nucleotide sequence that may lead to transcriptional silencing upon interaction with PRC2. Components of PRC2 complex in the brain are below average levels in comparison with many tissues. For this reason, we may conclude that the silencing potential is more characteristic of other tissues than the brain. Transcription variants 1 and 2 are expressed only in the brain. Moreover, truncated variant 3 with the deletion of Alu cluster has an expanded range of tissue expression.

The main part in the centre of the first long intronic RNA (*GABRB3,* variant1,2 and partially variant 3) can recruit RNA-binding serine-arginine proteins *SRSF1,2, PTB P* and/or NS. Other portions have potential binding sites for *hnRNP C, L, G*, and *YY1* binding proteins. This potential binding to different proteins is tissue-dependent, which should correspond with their concentration levels in the nucleus (Proteomics data). According to statistical data, the first long introns are subject to post-transcriptional splicing more frequently than others, and therefore, they can potentiate the creation of the elevated component levels of the future spliceosome as a whole in the *GABRB3* gene region up to the transcriptional termination. An area adjacent to the pre-mRNA 3’-end with a higher number of intron-exon alternations (*GABRB3,* variant1-4) is also enriched in serine-arginine protein *SRSF1,2* RNA-binding sites and strong isolated NP signals reducing the transcription rate. All of these reasons may explain, to some extent, the changes in processing efficiency in this region (processing slowdown and/or increasing accuracy of splicing).

An area in the long first intron adjacent to the pre-mRNA 5’-end (*GABRB3,* in variant 1,2 and partially in variant 3) is enriched in NP clusters, in *CTCF* binding sites, in cryptic polyadenylation sites, provoking transcription pausing and *SRSF2* binding sites in a tissue-specific manner that may facilitate the participation of *SRSF2* protein in RP-II pause release and, as a whole, in accelerating the transcription up to high rate of elongation characteristic of long introns. A similar situation was identified for an area adjacent to the pre-mRNA 5’-end (*GABRA5*).

For the 15q11-12 locus, the chromosome homologous somatic pairing in human genome was identified as a rare event, in contrast to a similar frequent phenomena characteristic of Diptera, for which such events are associated with the presence of numerous bidirectional transcripts at the sites of pairing. For locus 15q11-12, we also identified bidirectional transcripts in the GenBank as annotated ones as well as *in silico* predicted. Folding of long intron RNAs in pairs of corresponding bidirectional transcripts also identified some reproducible substructures with almost identical coordinates that may easily interact with each other by numerous motif annealing in ss loops for (+) and (-) strands, thus initiating the homologous pairing.

## Acknowledgements

We are grateful to Ryazanskii S.S. for technical assistance in computer service and providing technical support, Suzdaleva M.V. and Mikhailova K.B. for technical help.

---

**Fig. S1. Overall number and density of sites for RNA-binding proteins.**

Density of *SRSF1*-binding sites, fragments f1-f5 are determined as in Fig. 2. (B) Overall number of *SRSF1* binding sites. (C) Density of *SRSF2* binding sites. (D) Overall number of *SRSF2-* binding sites. (E) Density of *SRSF5-*binding sites. (F) Overall number of *SRSF5-*binding sites.

Fig. S2. Secondary structure of ncRNA Malat1 (EF177381).

Table S1. Nucleotide sequence localization of elements for the *Homo sapiens* locus 15q11-12.

## References

1. Hollander D, Naftelberg S, Lev-Maor G, Kornblihtt AR, Ast G. How Are Short Exons Flanked by Long Introns Defined and Committed to Splicing? Trends Genet. 2016; 32: 596–606.

2. Rosonina E, Blencowe BJ. Analysis of the requirement for RNA polymerase II CTD heptapeptide repeats in pre-mRNA splicing and 3’-end cleavage. RNA. 2004; 10: 581–589.

3. de la Mata M, Kornblihtt AR. RNA polymerase II C-terminal domain mediates regulation of alternative splicing by SRp20. Nat Struct Mol Biol. 2006; 13: 973–980.

4. Hsin JP, Manley JL. The RNA polymerase II CTD coordinates transcription and RNA processing. Genes Dev. 2012; 26: 2119–2137.

5. Bentley DL. Coupling mRNA processing with transcription in time and space. Nat Rev Genet. 2014; 15: 163–175.

6. Bird G, Zorio DA, Bentley DL RNA polymerase II carboxy-terminal domain phosphorylation is required for cotranscriptional pre-mRNA splicing and 3’-end formation. Mol Cell Biol. 2004; 24: 8963–8969.

7. Nojima T, Gomes T, Grosso ARF, Kimura H, Dye MJ, Dhir S, Carmo-Fonseca M, Proudfoot NJ. Mammalian NET-Seq Reveals Genome-wide Nascent Transcription Coupled to RNA Processing. Cell. 2015; 161: 526–540.

8. Sims RJ, Millhouse S, Chen CF, Lewis BA, Erdjument-Bromage H, Tempst P, Manley JL, Reinberg D. Recognition of trimethylated histone H3 lysine 4 facilitates the recruitment of transcription postinitiation factors and pre-mRNA splicing. Mol Cell. 2007; 28: 665–676.

9. Pray-Grant MG, Daniel JA, Schieltz D, Yates JR 3rd, Grant PA. Chd1 chromodomain links histone H3 methylation with SAGA-and SLIK-dependent acetylation. Nature. 2005; 433: 434– 438.

10. Martinez E, Palhan VB, Tjernberg A, Lymar ES, Gamper AM, Kundu TK, Chait BT, Roeder RG. Human STAGA complex is a chromatin-acetylating transcription coactivator that interacts with pre-mRNA splicing and DNA damage-binding factors in vivo. Mol. Cell. Biol. 2001; 20: 6782–6795.

11. Kfir N, Lev-Maor G, Glaich O, Alajem A, Datta A, Sze SK, et al. SF3B1 association with chromatin determines splicing outcomes. Cell Rep. 2015; 11: 618–629.

12. Batsché E^1^, Yaniv M, Muchardt C. Nat Struct Mol Biol. The human SWI/SNF subunit Brm is a regulator of alternative splicing. Nat Struct Mol Biol. 2006; 13: 22–29.

13. Gelfman S, Burstein D, Penn O, Savchenko A, Amit M, Schwartz S, et al. Genome Res. 2012;22(1):35-50. Changes in exon-intron structure during vertebrate evolution affect the splicing pattern of exons. Genome Res. 2012; 22: 35–50.

14. de la Mata M, Lafaille C, Kornblihtt AR. First come, first served revisited: factors affecting the same alternative splicing event have different effects on the relative rates of intron removal. RNA. 2010; 16: 904–912.

15. Bhatt DM, Pandya-Jones A, Tong AJ, Barozzi I, Lissner MM, Natoli G, et al. Transcript dynamics of proinflammatory genes revealed by sequence analysis of subcellular RNA fractions. Cell. 2012; 150: 279–290.

16. Huang S, Spector DL. Intron-dependent recruitment of pre-mRNA splicing factors to sites of transcription. J Cell Biol. 1996; 133: 719–732.

17. Kaida D, Berg MG, Younis I, Kasim M, Singh LN, Wan L, et al. U1 snRNP protects pre-mRNAs from premature cleavage and polyadenylation. Nature. 2010; 468: 664–668.

18. Oh JM, Di C, Venters CC, Guo J, Arai C, So BR, et al. U1 snRNP telescripting regulates a sizefunction-stratified human genome. Nat Struct Mol Biol. 2017; 24: 993–999.

19. Bernard D, Prasanth KV, Tripathi V, Colasse S, Nakamura T, Xuan Z, et al. A long nuclearretained noncoding RNA regulates synaptogenesis by modulating gene expression. EMBO J. 2010; 29: 3082–3093.

20. Melcák I, Cermanová S, Jirsová K, Koberna K, MalínsKý J, Raska I. Nuclear pre-mRNA compartmentalization: trafficking of released transcripts to splicing factor reservoirs. Mol Biol Cell. 2000; 11: 497–510.

21. Spector DL, Lamond AI. Nuclear speckles. Cold Spring Harb Perspect Biol. 2011; 3(2). doi:10.1101/cshperspect.a000646.

22. Hall LL, Smith KP, Byron M, Lawrence JB. Molecular anatomy of a speckle. Anat Rec A Discov Mol Cell Evol Biol. 2006; 288: 664–675.

23. Shopland LS, Johnson CV, Lawrence JB. Evidence that all SC-35 domains contain mRNAs and that transcripts can be structurally constrained within these domains. J Struct Biol. 2002; 140: 131–139.

24. Cmarko D, Verschure PJ, Martin TE, Dahmus ME, Krause S, Fu XD, et al. Ultrastructural analysis of transcription and splicing in the cell nucleus after bromo-UTP microinjection. Mol Biol Cell. 1999; 10: 211–223.

25. Fakan S. Perichromatin fibrils are in situ forms of nascent transcripts. Trends Cell Biol. 1994; 4: 86–90.

26. Query CC, McCaw PS, Sharp PA. A minimal spliceosomal complex A recognizes the branch site and polypyrimidine tract. Mol Cell Biol. 1997; 17: 2944–53.

27. Melcak I, Melcakova S, Kopsky V, Vecerova J, and Raska I. Prespliceosomal assembly on microinjected precursor mRNA takes place in Nuclear Speckles. Molecular Biology of the Cell. 2001; 12: 393–406.

28. Wada Y, Ohta Y, Xu M, Tsutsumi S, Minami T, Inoue K, et al. A wave of nascent transcription on activated human genes. Proc Natl Acad Sci U S A. 2009; 106: 18357–18361.

29. LaSalle JM, Lalande M. Homologous association of oppositely imprinted chromosomal domains. Science. 1996; 272: 725–728.

30. Thatcher KN, Peddada S, Yasui DH, Lasalle JM. Homologous pairing of 15q11-13 imprinted domains in brain is developmentally regulated but deficient in Rett and autism samples. Hum Mol Genet. 2005; 14: 785–797.

31. Hogart A, Nagarajan RP, Patzel KA, Yasui DH, Lasalle JM. 15q11-13 GABAA receptor genes are normally biallelically expressed in brain yet are subject to epigenetic dysregulation in autismspectrum disorders. Hum Mol Genet. 2007; 16: 691–703.

32. Husker J, Samuelssion T. and Strub K. Useful ‘junk’ Alu RNA in the human transcriptome Cell Moll. Life. Sci. 2007; 64: 1793–1800.

33. Lonsdale J, Thomas J, Salvatore M, Phillips R, Lo E, Shad S, et al. The Genotype-Tissue Expression (GTEx) project. GTEx Consortium. Nat Genet. 2013; 45: 580–585.

34. Wagstaff J, Knoll JH, Fleming J, Kirkness EF, Martin-Gallardo A, Greenberg F et al. Localization of the gene encoding the GABAA receptor beta 3 subunit to the Angelman/Prader-Willi region of human chromosome 15. Am J Hum Genet. 1991; 49: 330–337.

35. Magenis RE, Toth-Fejel S, Allen LJ, Black M, Brown MG, Budden S, et al. Comparison of the 15q deletions in Prader-Willi and Angelman syndromes: specific regions, extent of deletions, parental origin, and clinical consequences. Am J Med Genet. 1990; 35: 333–349.

36. Lalande M, Minassian BA, DeLorey TM, Olsen RW. Parental imprinting and Angelman syndrome. Adv Neurol. 1999; 79: 421–429.

37. Neubert G, von Au K, Drossel K, Tzschach A, Horn D, Nickel R, et al. Angelman syndrome and severe infections in a patient with de novo 15q11.2-q13.1 deletion and maternally inherited 2q21.3 microdeletion. Gene. 2013; 512: 453–455.

38. Jiang YH, Pan Y, Zhu L, Landa L, Yoo J, Spencer C, et al. Altered ultrasonic vocalization and impaired learning and memory in Angelman syndrome mouse model with a large maternal deletion from Ube3a to Gabrb3. PLOS One. 2010; 20; 5(8):e12278.

39. Francke U. Imprinted genes in the Prader-Willi deletion. Novartis Found Symp. 1998; 214: 264–275.

40. Thatcher KN, Peddada S, Yasui DH, Lasalle JM. Homologous pairing of 15q11-13 imprinted domains in brain is developmentally regulated but deficient in Rett and autism samples. Hum Mol Genet. 2005; 14: 785–97.

41. Hodges LM, Fyer AJ, Weissman MM, Logue MW, Haghighi F, Evgrafov O et al. Evidence for linkage and association of GABRB3 and GABRA5 to panic disorder. Neuropsychopharmacology. 2014; 39: 2423–2431.

42. Markham, N. R. & Zuker, M. UNAFold: software for nucleic acid folding and hybridization. 2008; In Keith, J. M., editor, Bioinformatics, Volume II. Structure, Function and Applications, number 453 in Methods in Molecular Biology, chapter 1, pages 3–31. Humana Press, Totowa, NJ. ISBN 978-1-60327-428-9.

43. Zuker M. Mfold web server for nucleic acid folding and hybridization prediction. Nucleic Acids Res. 2003; 31: 3406–3415.

44. Fedoseyeva V, Zharinova I, Alexandrov AA. Secondary structure-stretched forms of long intron RNA products from the view point of initiation of chromosome homologs somatic pairing. J Biomol Struct Dyn. 2015; 33: 869–876.

45. König J, Zarnack K, Rot G, Curk T, Kayikci M, Zupan B, et al. iCLIP reveals the function of hnRNP particles in splicing at individual nucleotide resolution. Nat Struct Mol Biol. 2010; 17: 909–915.

46. Burd CG and Dreyfuss G. RNA binding specificity of hnRNP A1: significance of hnRNP A1 high-affinity binding sites in pre-mRNA splicing. EMBO J. 1994; 13: 1197–1204.

47. Liu HX,, Zhang M, Krainer AR. Identification of functional exonic splicing enhancer motifs recognized by individual SR proteins. Genes Dev. 1998; 12: 1998–2012.

48. Cavaloc Y, Bourgeois CF, Kister L, and Stévenin J. The splicing factors 9G8 and SRp20 transactivate splicing through different and specific enhancers. RNA. 1999; 5: 468–483.

49. Liu HX, Chew SL, Cartegni L, Zhang MQ, Krainer AR. Exonic splicing enhancer motif recognized by human SC35 under splicing conditions. Mol Cell Biol. 2000; 20: 1063–1071.

50. Sikora D, Zhang D, Bojic T, Beeharry Y, Tanara A, Pelchat M. Identification of a binding site for ASF/SF2 on an RNA fragment derived from the hepatitis delta virus genome. PLOS One. 2013; 8(1):e54832.

51. Tacke R, Manley JL. The human splicing factors ASF/SF2 and SC35 possess distinct, functionally significant RNA binding specificities. EMBO J. 1995; 14: 3540–3551.

52. Wagner EJ, Garcia-Blanco MA. Polypyrimidine tract binding protein antagonizes exon definition. Mol Cell Biol. 2001; 21: 3281–3288.

53. Le Guiner C, Plet A, Galiana D, Gesnel MC, Del Gatto-Konczak F, Breathnach R. Polypyrimidine tract-binding protein represses splicing of a fibroblast growth factor receptor-2 gene alternative exon through exon sequences. J Biol Chem. 2001; 276: 43677–43687.

54. Spellman R, Rideau A, Matlin A, Gooding C, Robinson F, McGlincy N, et al. Regulation of alternative splicing by PTB and associated factors. Biochem Soc Trans. 2005; 33(Pt 3): 457–60.

55. Hertel KJ and Graveley BR. RS domains contact the pre-mRNA throughout spliceosome assembly. Trends Biochem Sci. 2005; 30: 115–118.

56. Shen H and Green MR. A pathway of sequential arginine-serine-rich domain-splicing signal interactions during mammalian spliceosome assembly. Mol Cell. 2004; 16: 363–373.

57. Shen H and Green MR. RS domains contact splicing signals and promote splicing by a common mechanism in yeast through humans. Genes Dev. 2006; 20: 1755–1765.

58. Staknis D and Reed R. SR proteins promote the first specific recognition of Pre-mRNA and are present together with the U1 small nuclear ribonucleoprotein particle in a general splicing enhancer complex. Mol Cell Biol. 1994; 14: 7670–7682.

59. Kanopka A, Muhlemann O., Akusjarvi G. Inhibition by SR proteins of splicing of a regulated adenovirus pre-mRNA. Nature. 1996; 381: 535– 538.

60. Cook CR and McNally MT. SR protein and snRNP requirements for assembly of the Rous sarcoma virus negative regulator of splicing complex in vitro. Virology. 1998; 42: 211–220.

61. Shen M, Mattox W. Activation and repression functions of an SR splicing regulator depend on exonic versus intronic-binding position. Nucleic Acids Res. 2012; 40: 428–437.

62. Bjork P, Jin S, Zhao J, Singh OP, Persson JO, Hellman U, et al. Specific combinations of SR proteins associate with single pre-messenger RNAs in vivo and contribute different functions. J Cell Biol. 2009; 184: 555–568.

63. Misteli T, Spector DL. RNA polymerase II targets pre-mRNA splicing factors to transcription sites in vivo. Mol Cell. 1999; 3: 697–705.

64. Tripathi V1, Song DY, Zong X, Shevtsov SP, Hearn S, Fu XD, et al. SRSF1 regulates the assembly of pre-mRNA processing factors in nuclear speckles. Mol Biol Cell. 2012; 23: 3694– 3706.

65. Bourgeois CF, Lejeune F, Stévenin J. Broad specificity of SR (serine/arginine) proteins in the regulation of alternative splicing of pre-messenger RNA. Prog Nucleic Acid Res Mol Biol. 2004; 78: 37–88.

66. Das S and Krainer AR. Emerging functions of SRSF1, splicing factor and oncoprotein, in RNA metabolism and cancer. Mol Cancer Res. 2014; 12: 1195–1204.

67. Graveley BR. Sorting out the complexity of SR protein functions. RNA. 2000; 6(9):1197–211.

68. Sanford JR, Ellis J, Cáceres JF. Multiple roles of arginine/serine-rich splicing factors in RNA processing. Biochem Soc Trans. 2005; 33(Pt 3): 443-6.

69. Fagerberg L, Hallström BM, Oksvold P, Kampf C, Djureinovic D, Odeberg J et al. Analysis of the human tissue-specific expression by genome-wide integration of transcriptomics and antibody-based proteomics. Mol Cell Proteomics. 2014; 13: 397–406.

70. Ji X, Zhou Y, Pandit S, Huang J, Li H, Lin CY, et al. SR proteins collaborate with 7SK and promoterassociated nascent RNA to release paused polymerase. Cell. 2013; 153: 855–868.

71. Lin S, Coutinho-Mansfield G, Wang D, Pandit S, Fu XD. The splicing factor SC35 has an active role in transcriptional elongation. Nat Struct Mol Biol. 2008; 15: 819–826.

72. Zhang J, Lieu YK, Ali AM, Penson A, Reggio KS, Rabadan R, et al. Disease-associated mutation in SRSF2 misregulates splicing by altering RNA-binding affinities. Proc Natl Acad Sci U S A. 2015; 112: E4726–4734.

73. Tyson-Capper AJ, Bailey J, Krainer AR, Robson SC, Europe-Finner GN. The switch in alternative splicing of cyclic AMP-response element modulator protein CREM{tau}2{alpha} (activator) to CREM{alpha} (repressor) in human myometrial cells is mediated by SRp40. J Biol Chem. 2005; 280: 34521–34529.

74. Romanelli MG, Diani E, Lievens PM. New insights into functional roles of the polypyrimidine tract-binding protein. Int J Mol Sci. 2013; 14: 22906–22932.

75. Lamichhane R, Daubner GM, Thomas-Crusells J, Auweter SD, Manatschal C, Austin KS, et al. RNA looping by PTB: Evidence using FRET and NMR spectroscopy for a role in splicing repression. Proc Natl Acad Sci U S A. 2010; 107: 4105–4110.

76. Matlin AJ, Southby J, Gooding C, Smith CW. Repression of alpha-actinin SM exon splicing by assisted binding of PTB to the polypyrimidine tract. RNA. 2007; 13: 1214–1223.

77. Saulière J, Sureau A, Expert-Bezançon A, Marie J. The polypyrimidine tract binding protein (PTB) represses splicing of exon 6B from the beta-tropomyosin pre-mRNA by directly interfering with the binding of the U2AF65 subunit. Mol Cell Biol. 2006; 26: 8755–8769.

78. Ashiya M, Grabowski PJ. A neuron-specific splicing switch mediated by an array of pre-mRNA repressor sites: evidence of a regulatory role for the polypyrimidine tract binding protein and a brain-specific PTB counterpart. RNA. 1997; 3: 996–1015.

79. Simpson PJ, Monie TP, Szendroi A, Davydova N, Tyzack JK, Conte MR, et al. Structure and RNA interactions of the N-terminal RRM domains of PTB. Structure. 2004; 12: 1631–43.

80. Petoukhov MV, Monie TP, Allain FH, Matthews S, Curry S, Svergun DI. Conformation of polypyrimidine tract binding protein in solution. Structure. 2006; 14: 1021–1027.

81. Clerte C, Hall KB. Characterization of multimeric complexes formed by the human PTB1 protein on RNA. RNA. 2006; 12: 457–475.

82. Hui J, Stangl K, Lane WS, Bindereif A. HnRNP L stimulates splicing of the eNOS gene by binding to variable-length CA repeats. Nat Struct Biol. 2003; 10: 33–37.

83. Hui J, Hung L-H, Heiner M, Schreiner S, Neumuller N, Reither G et al. Intronic CA-repeat and CA-rich elements: a new class of regulators of mammalian alternative splicing. EMBO Journal. 2005; 24: 1988–1998.

84. Heinrich B, Zhang Z, Raitskin O, Hiller M, Benderska N, Hartmann AM, et al. Heterogeneous nuclear ribonucleoprotein G regulates splice site selection by binding to CC(A/C)-rich regions in pre-mRNA. J Biol Chem. 2009; 284: 14303–14315.

85. Burd CG, Dreyfuss G. Conserved structures and diversity of functions of RNA-binding proteins. Science. 1994; 265: 615–621.

86. Expert-Bezançon A, Sureau A, Durosay P, Salesse R, Groeneveld H, Lecaer JP, et al. hnRNP A1 and the SR proteins ASF/SF2 and SC35 have antagonistic functions in splicing of beta-tropomyosin exon 6B. J Biol Chem. 2004; 279: 38249–38259.

87. Liu X, Ishizuka T, Bao HL, Wada K, Takeda Y, Iida K, et al. Structure-Dependent Binding of hnRNPA1 to Telomere RNA. J Am Chem Soc. 2017; 139: 7533–7539.

88. Krüger AC, Raarup MK, Nielsen MM, Kristensen M, Besenbacher F, Kjems J, Birkedal V. Interaction of hnRNP A1 with telomere DNA G-quadruplex structures studied at the single molecule level. Eur Biophys J. 2010; 39: 1343:1350.

89. Munroe SH, Dong XF. Heterogeneous nuclear ribonucleoprotein A1 catalyzes RNA.RNA annealing. Proc Natl Acad Sci U S A. 1992; 89: 895–899.

90. Ishihama Y, Tadakuma H, Tani T, Funatsu T. The dynamics of pre-mRNAs and poly(A)+ RNA at speckles in living cells revealed by iFRAP studies. Exp Cell Res. 2008; 314: 748–762.

91. Wang J, Cao LG, Wang YL, and Pederson T. Localization of pre-messenger RNA at discrete nuclear sites. Proc Natl Acad Sci U S A. 1991; 88: 7391–7395.

92. Engreitz JM, Sirokman K, McDonel P, Shishkin AA, Surka C, Russell P, et al. RNA-RNA interactions enable specific targeting of noncoding RNAs to nascent Pre-mRNAs and chromatin sites. Cell. 2014; 159: 188–199.

93. Reid DC, Chang BL, Gunderson SI, Alpert L, Thompson WA, Fairbrother WG. Next-generation SELEX identifies sequence and structural determinants of splicing factor binding in human pre-mRNA sequence. RNA. 2009; 15: 2385–2397.

94. Engreitz JM, Sirokman K, McDonel P, Shishkin AA, Surka C, Russell P, et al. RNA-RNA interactions enable specific targeting of noncoding RNAs to nascent Pre-mRNAs and chromatin sites. Cell. 2014; 159: 188–199.

95. Athanasiadis A, Rich A, Maas S. Widespread A-to-I RNA editing of Alu-containing mRNAs in the human transcriptome. PLOS Biol. 2004; 2(12):e391.

96. Maenner S, Blaud M, Fouillen L, Savoye A, Marchand V, Dubois A, et al. 2-D structure of the A region of Xist RNA and its implication for PRC2 association. PLOS Biol. 2010; 8(1):e1000276.

97. Zhao J, Sun BK, Erwin JA, Song J-J, and Lee JT. Polycomb proteins targeted by a short repeat RNA to the mouse X-chromosome Science. 2008; 322: 750–756.

98. Wutz A, Rasmussen TP, Jaenisch R. Nat Genet. Chromosomal silencing and localization are mediated by different domains of Xist RNA. Nat Genet. 2002; 30: 167–74.

99. Davidovich C, Wang X, Cifuentes-Rojas C, Goodrich KJ, Gooding AR, Lee JT, et al. Toward a consensus on the binding specificity and promiscuity ofPRC2 for RNA. Mol Cell. 2015; 57: 552–558.

100. Kanhere A, Viiri K, Araújo CC, Rasaiyaah J, Bouwman RD, Whyte WA, et al. Short RNAs are transcribed from repressed polycomb target genes and interact with polycomb repressive complex-2. Mol Cell. 2010; 38: 675–688.

101. Pham CD, He X, Schnitzler GR. Divergent human remodeling complexes remove nucleosomes from strong positioning sequences. Nucleic Acids Res. 2010; 38: 400–413.

102. Fan HY, Trotter KW, Archer TK, Kingston RE. Swapping function of two chromatin remodeling complexes. Mol Cell. 2005; 17: 805–815.

103. Govind CK, Zhang F, Qiu H, Hofmeyer K, Hinnebusch AG. Gcn5 promotes acetylation, eviction, and methylation of nucleosomes in transcribed coding regions. Mol Cell. 2007; 25:31–42.

104. Kassabov SR, Zhang B, Persinger J, Bartholomew B. SWI/SNF unwraps, slides, and rewraps the nucleosome. Mol Cell. 2003; 11: 391–403.

105. Samara NL, Wolberger C. A new chapter in the transcription SAGA. Curr Opin Struct Biol. 2011; 21: 767–774.

106. Schwartz S, Meshorer E, Ast G. Chromatin organization marks exon-intron structure. Nat Struct Mol Biol. 2009; 16: 990–995.

107. Rybak-Wolf A, Stottmeister C, Glažar P., MarvinJens M, Pino N, Giusti S, et al. Circular RNAs in the Mammalian Brain Are Highly Abundant, Conserved, and Dynamically Expressed. Mol Cell 2015; 58: 870–885. http://www.circbase.org/cgi-bin/simplesearch.cgi

108. Joyce EF, Williams BR, Xie T, and Wu CT. Identification of genes that promote or antagonize somatic homolog pairing using a high-throughput FISH–based screen. PLOS Genetics. 2012; 8(5):e1002667.

109. Krueger C, King MR, Krueger F, Branco MR, Osborne CS, Niakan KK, et al. Pairing of homologous regions in the mouse genome is associated with transcription but not imprinting status. PLOS One. 2012; 7(7):e38983.

110. Apte MS, Meller VH. Homologue pairing in flies and mammals: gene regulation when two are involved. Genet Res Int. 2012; 2012:430587, 9 pages.

